# Interpretable compositional computation with recurrent neural networks

**DOI:** 10.64898/2026.06.23.733979

**Authors:** Louis Pezon, Alexander van Meegen

## Abstract

Flexible cognition utilizes reusable components to enable rapid adaptation of behavior to different contexts or tasks. Analysis of artificial neural networks trained on multiple tasks suggested that this compositionality is supported by dynamical structures which are shared and re-used across tasks. However, the nature of these shared components, and how they can be used in a task-dependent manner, remained unclear. Here, we develop a theory of interpretable compositional computation based on shared dynamical structures in the low-dimensional latent space of low-rank recurrent neural networks. We show that these *shared latent components* are not immediately visible in the neural activity, and are thus compatible with task-dependent activity. We identify hallmarks of shared latent components both in the connectivity statistics and the neural representations. These hallmarks yield testable predictions for the network’s response to specific perturbation experiments. Finally, we identify distinct loci where task-dependence can enter the computation, allowing us to characterize qualitatively different solutions to compositional tasks. In summary, our theory provides a mechanistic understanding and testable hallmarks of compositional computation via shared components in low-rank networks.

## I. INTRODUCTION

A key feature of human cognition is the ability to flexibly re-use learned concepts and behaviors [1]. For example, once a trumpeter masters air flow and harmonics, she can use these skills across a vast repertoire of musical pieces. Furthermore, if she is faced with a new instrument, such as an Alphorn, she can quickly adapt using her existing repertoire of skills. However, compositionality—exemplified by the player flexibly combining her learned skills—is a hard task for artificial neural networks [2–5], including state-of-the-art transformer-based networks [6–8]. While compositionality alone poses a significant challenge, there is an additional layer of complexity: natural behavior is inherently dynamic, which needs to be reflected by the computation [9–11]. How do the neural circuits in the brain implement the dynamic and compositional computations which enable the players performance?

Finding appropriate models for dynamic and compositional computation is an ongoing challenge. One prominent approach is to train artificial networks on a family of tasks with a common structure [12–15]. An analysis of the trained networks revealed that the network computation can reflect this structure: the trained networks re-purposed representations [12] or “dynamical motifs” [14, 15] across similar tasks — a finding that parallels observations in prefrontal cortex [16]. In these cases, the networks learned to implement a compositional computation using “shared components”.

However, the nature of these components is not clear. Even if neural representations or dynamics involve shared subspaces, neural activity remains task-dependent [15–17]. Furthermore, a compositional multi-task computation should combine shared components with task-dependent ones. Consider, for example, a family of tasks that all require memorizing an indicated direction. Such a family of tasks can be solved using a ring attractor [18–21] that is shared across tasks [15]. Yet, a shared ring attractor alone cannot perform task-specific computations: it must be used in a task-dependent manner. This raises two questions: First, how can shared components be reconciled with task-dependent activity? And second, how can they be used in a task-dependent manner to enable flexible computation in a single network?

To address these questions, we assume that the computations are implemented by recurrent neural network dynamics [9]. We exploit the fact that the task structure of typical neuroscience experiments is low-dimensional [22, 23], which means that the tasks can be solved using low-dimensional dynamics. Indeed, understanding the computation of a neural population in terms of low-dimensional dynamics has been a striking success story [10, 11, 17, 24–33]. Such low-dimensional dynamics are, in turn, naturally implemented in low-rank networks [27].

Here, we build on the extensive theory of low-rank networks [27, 31, 32, 34–45] to understand how a network with a generic structure can solve multiple tasks by combining shared and task-dependent components. To this end, we develop a theory of compositional computation in low-rank networks. A direct result of the theory is that shared components are not immediately obvious in neural activity, and thus consistent with task-dependent activity. Nonetheless, we show that computation via shared components leaves testable hallmarks in the connectivity statistics. Furthermore, shared components impose specific constraints on the geometry of neural activity across tasks, for which we propose a test based on perturbation experiments. Next, we identify two loci where each task variable can enter a computation via shared components, that can be identified via perturbation experiments. Generalizing our framework to a compositional family of tasks [16], we build a systematic catalog of qualitatively different solutions to the same tasks. Finally, we construct explicit examples of how shared components can enable solving complex compositional tasks.

## II. RESULTS

### A. Multi-task computation in low-rank RNNs

We develop a theory of multi-task computation via shared computational components (or “motifs” [15]) in recurrent neural networks (RNNs). To this end, we consider artificial neural networks consisting of *N* neurons, characterized by their firing rates ***r***(*t*) = (*r*_1_(*t*), …, *r*_*N*_ (*t*)). The vector of firing rates ***r***(*t*), called the population activity, fully describes the instantaneous state of the network. We assume that the same RNN is used across different tasks and that the task identity has been correctly inferred from the context, for example by a separate network that is not part of our model. During a trial, the task identity is directly provided to the network, encoded as a constant input ***c***; in addition, the network receives a time-dependent stimulus ***s***(*t*) (Fig. 1A).

**Figure 1.**
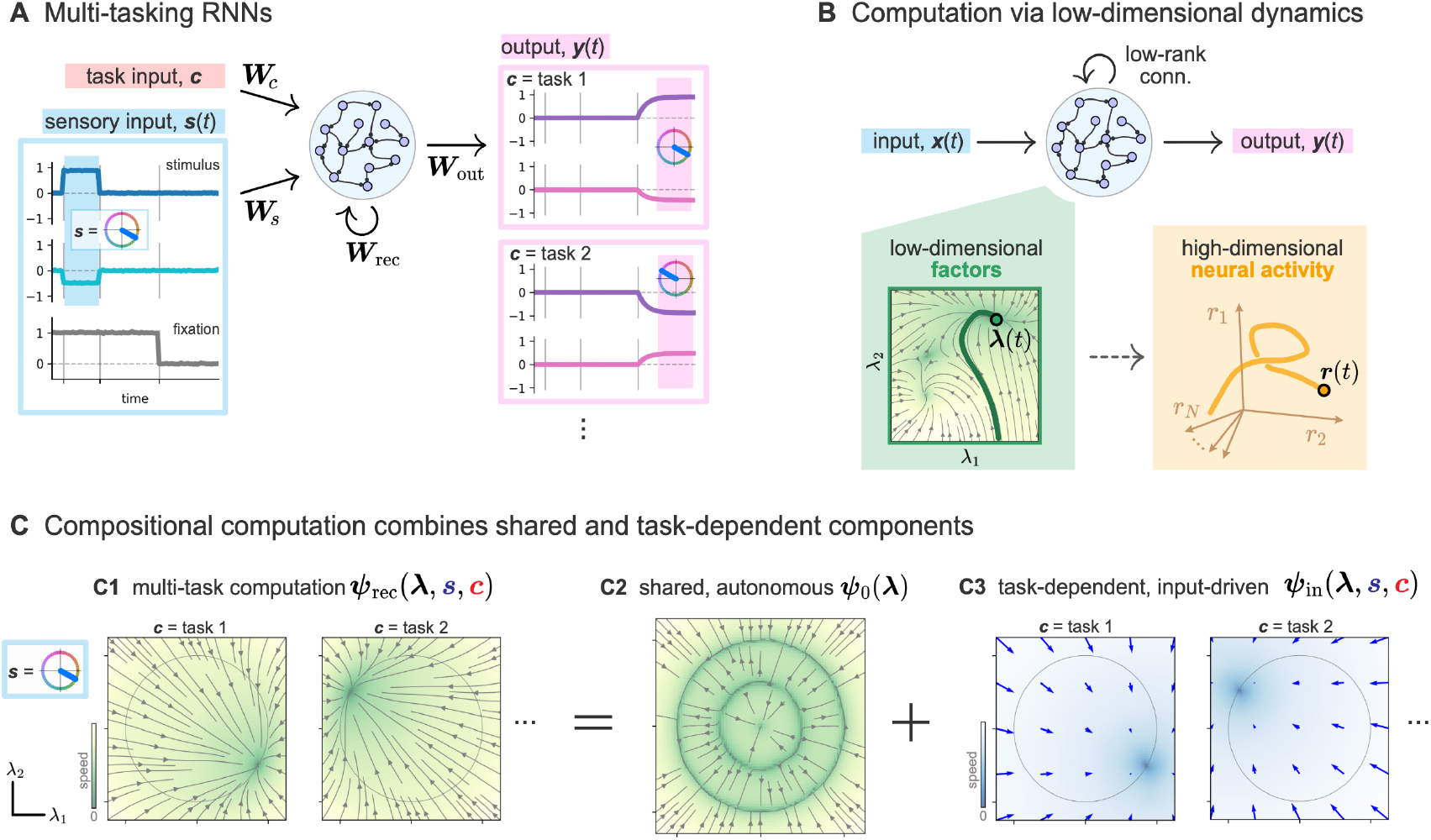
Compositional computation using shared components in low-rank RNNs. **A.** Multi-tasking RNNs receive a stimulus ***s***(*t*) and a constant input ***c*** encoding task identity. The transformation from stimulus ***s*** (left; blue direction on the circle represented by *s*_1_ = cos *α, s*_2_ = sin *α*) to network output ***y*** depends on the task identity (right). **B**. Low-rank RNNs naturally implement low-dimensional latent dynamics ***λ***(*t*) in their high-dimensional neural activity ***r***(*t*). These latent dynamics fully determine the input-output transformation and the neural activity. **C**. In multi-tasking low-rank RNNs, the autonomous part ***ψ***_0_(***λ***) of the low-dimensional recurrent dynamics ***ψ***_rec_(***λ, s, c***) can serve as a shared component which is re-used across tasks. The combination with task-dependent input-driven dynamics and readout enables compositional computation.

The dynamics of the RNN are

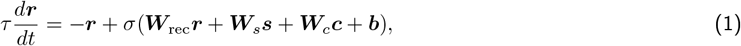

where ***W***_rec_ denotes the recurrent connectivity, ***W***_*s*_ and ***W***_*c*_ the input weights from stimulus ***s*** and task input ***c***, respectively, ***b*** the neuron-specific bias, *τ* the time constant of the dynamics, and *σ* the neuronal nonlinearity (softplus for all simulations). The final output is a linear readout of the rates,

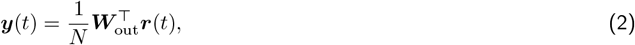

where ***W***_out_ denotes the readout weights.

RNNs are capable of performing computations via low-dimensional latent dynamics, embedded in the high-dimensional space of firing rates ***r***(*t*) (Fig. 1B, [10, 17, 24–27, 29–32]). Low-rank connectivity naturally gives rise to such dynamics [27, 31, 32, 34–45]. Following this prominent line of work, we assume that the connectivity is low-rank, 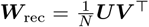 where ***U*** and ***V*** are *N* ×*P* matrices with *P* ≪ *N*. This allows us to identify a small number *P* of factors, 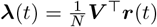, and introduce the recurrent flow-field ***ψ***_rec_ that governs their dynamics:

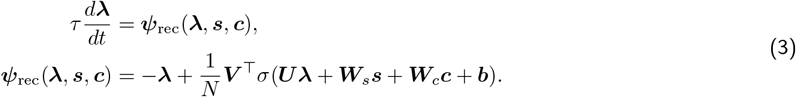

The trajectory of the factors ***λ***(*t*), together with the inputs, ***s***(*t*) and ***c***, fully determine the outputs as

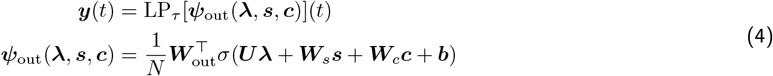

where the readout flow-field ***ψ***_out_(***λ, s, c***) controls the nonlinear transformation from factors to outputs, and LP_*τ*_ denotes low-pass filtering^1^ with time constant *τ*. It follows from Eqs. (3)-(4) that the dynamics and output of the network—and thus, the computation it performs—is entirely determined by the low-dimensional flow-fields ***ψ***_rec_ and ***ψ***_out_.

It is convenient to further distinguish the autonomous dynamics, which implement computation in the absence of stimulus, for example during a memory period, from the input-driven dynamics that mediate the effect of the stimulus. This corresponds to the unique decomposition of the recurrent flow-field into two terms [33]:

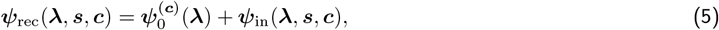

where 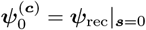 describes the latent dynamics for task ***c*** in the absence of stimulus (i.e., ***s*** = 0), and the input-driven flow-field ***ψ***_in_ satisfies ***ψ***_in ***s***=0_ = 0. The decomposition of Eq. (5) allows us to express how a network can combine shared and task-dependent components: we propose that *the autonomous dynamics provide the shared components*. Formally, we require that autonomous dynamics are the same across tasks and do not depend on the task input ***c***, such that one can write

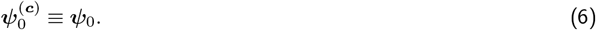

To implement task-dependent computation, the task identity ***c*** can modulate both the effect of the stimulus via the input-driven flow-field ***ψ***_in_, and the network’s output via ***ψ***_out_ (Fig. 1C).

As an illustration, we consider a set of two stylized memory tasks [15]: the *pro* and *anti* memory tasks. Both tasks require memorizing a direction indicated by a transient input. After a delay, either the stimulus angle (‘pro’ task), or its opposite (‘anti’ task), should be reported (Fig. 1A). These tasks can be solved using a shared latent ring attractor (Fig. 1C2), combined with a task-dependent input-driven flow-field: depending on the task, either the stimulus angle, or its opposite, is stored using the ring attractor (Fig. 1C3). Here, the task dependence is resolved by the interaction of the input with the ring attractor. Afterwards, a simple readout of the stored angle is sufficient to solve both tasks.

A key advantage of considering the low-dimensional flow-field ***ψ***_rec_ is that it provides a highly interpretable perspective on the latent dynamics. For example, it was recently shown that evidence accumulation is best understood in terms of a combination of input-driven dynamics for evidence accumulation and autonomous dynamics to fix the decision [33]. In the context of multi-task networks, the flow-field supports interpretability by explicitly separating shared and task-dependent components. For now, we consider for simplicity the case where the network uses the same shared autonomous dynamics across all tasks. However, our results directly generalize to the case when only subsets of tasks in a family use shared components, as we will show later.

### B. Shared components coexist with task-dependent neural activity

Shared components across tasks seem to be at odds with the observation that neural activity often evolves in different subspaces across contexts or tasks (Fig. 2A, [17, 24, 25, 30, 46–51]). Perhaps counter-intuitively, we show that there is no such contradiction because the shared components are on the level of the factors. Going from the factors ***λ***(*t*) to the population activity ***r***(*t*) reveals that shared latent components can coexist with task-dependent activity. Indeed, the vector of firing rates,

**Figure 2.**
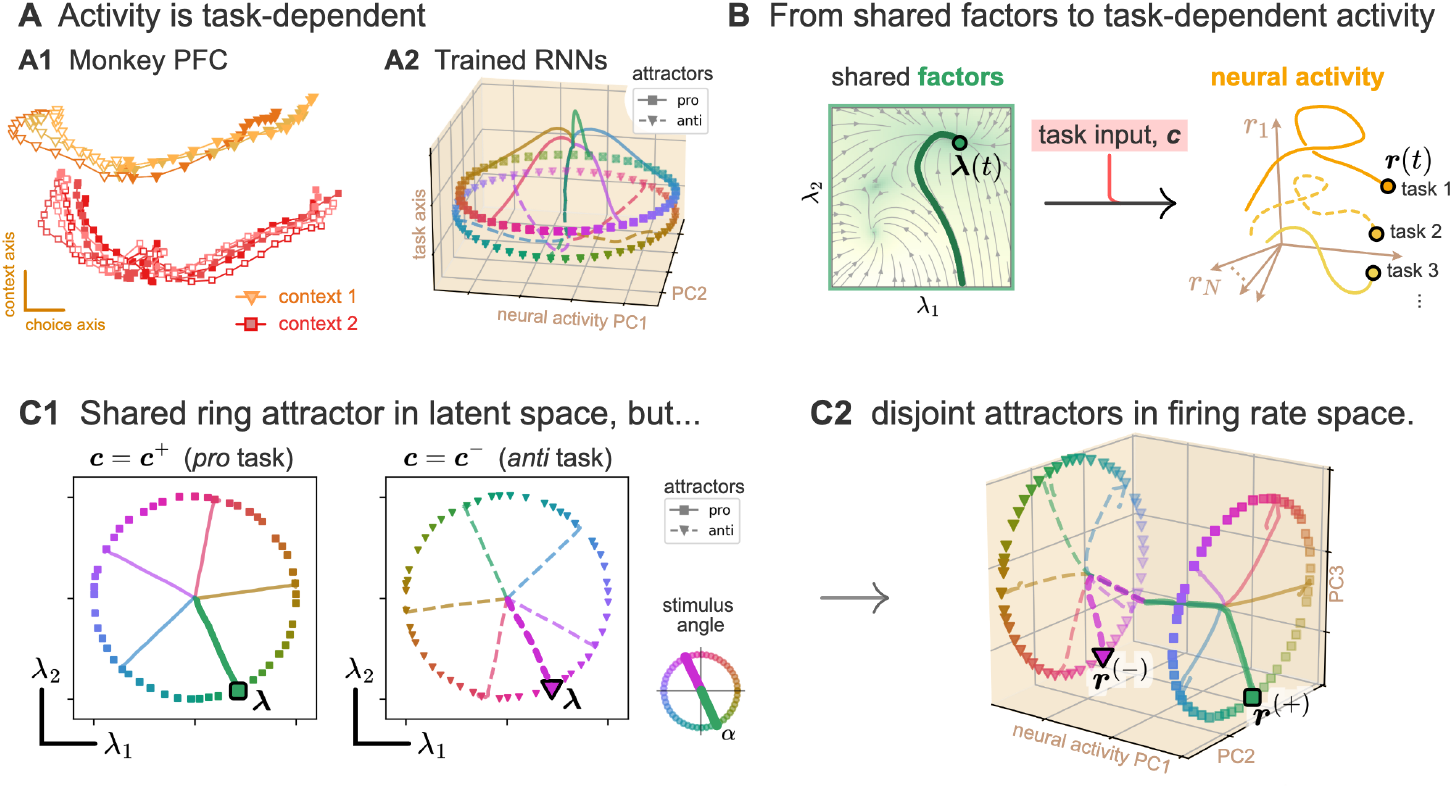
Computation via shared latent components is consistent with task-dependent neural activity. **A.** Neural activity is task-dependent in monkey prefrontal cortex during context-dependent decision making (A1; data from Mante et al. [17]) and in multitasking RNNs (A2; data from Driscoll et al. [15]). **B**. Task-shared dynamics in latent space do not imply task-shared neural activity, since the activity depends on the task input ***c*. C**. Shared latent ring attractors for memory pro/anti tasks (C1) lead to disjoint attractors in firing rate space (C2; first three principal components shown). For a given stimulus angle *α* in the ‘pro’ task (left), or its opposite in the ‘anti’ task (right), factors ***λ*** follow the same trajectory (C1) while firing rates are separated across tasks (C2). In panel A1, projection axes were obtained using TDR [17]; activity is trial-averaged over motion conditions; and trajectories are styled according to context (colors and marker shapes), and motion direction (filled / empty markers) and coherence (color shades). In panels A2 and C2, PCA was performed on neural activity during the memory period. In A2, the ‘task’ axis was obtained by training a linear classifier to separate the two rings.

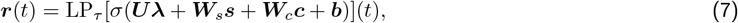

depends on both the factors ***λ*** as well as the task input ***c***. Accordingly, even if the factors ***λ*** follow identical dynamics across different tasks, the neural activity can still be task-dependent (Fig. 2B).

To illustrate this point, we consider again the pro/anti memory tasks and their solution using a shared ring attractor (Fig. 1C). We created a low-rank network that implements this solution in its latent dynamics (Methods). In the latent space, the same ring attractor is reused for both pro- and anti-task (Fig. 2C1). However, as shown above, the shared ring attractor in the latent space does not imply a shared ring attractor in the space of the neurons’ firing rates ***r***. Accordingly, the representations of the memorized stimulus form two disjoint rings, one for each task (Fig. 2C2), such that task identity is straightforward to decode from the first principal components of the activity.

The coexistence of shared latent components and task-dependent neural activity can be interpreted geometrically: the task-dependent part of the activity takes place in the nullspace of ***V*** (i.e., orthogonal to its column vectors), such that it does not affect the factors 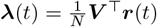. More precisely, the projection of the task-dependent activity on ***V*** needs to be negligible; strict confinement to the nullspace is not required. Since ***V*** spans only a *P* ≪ *N* dimensional subspace, there is ample room for task-dependent activity. Thus, the possibility of task-dependent activity despite shared latent components is a generic feature in low-rank networks, by no means restricted to the illustrative example used above. Conversely, and consistent with experimental evidence [16, 17], shared latent components imply that there is at least a *P*-dimensional subspace in which the activity is shared across tasks, which is the subspace spanned by ***V***.

### C. Shared components shape network connectivity

Computation via shared latent dynamics is compatible with task-dependent neural activity. Yet, a given family of tasks typically does not admit a unique compositional solution. A RNN could in principle solve each task using specific dynamics, without reusing components across tasks. For instance, the two ‘pro’ and ‘anti’ tasks presented above could be solved using one latent ring attractors for each task, rather than a shared one (Fig. 3A). What, then, distinguishes these different types of solution? As an answer, we identified specific testable constraints on the low-rank connectivity statistics imposed by shared autonomous latent dynamics.

**Figure 3.**
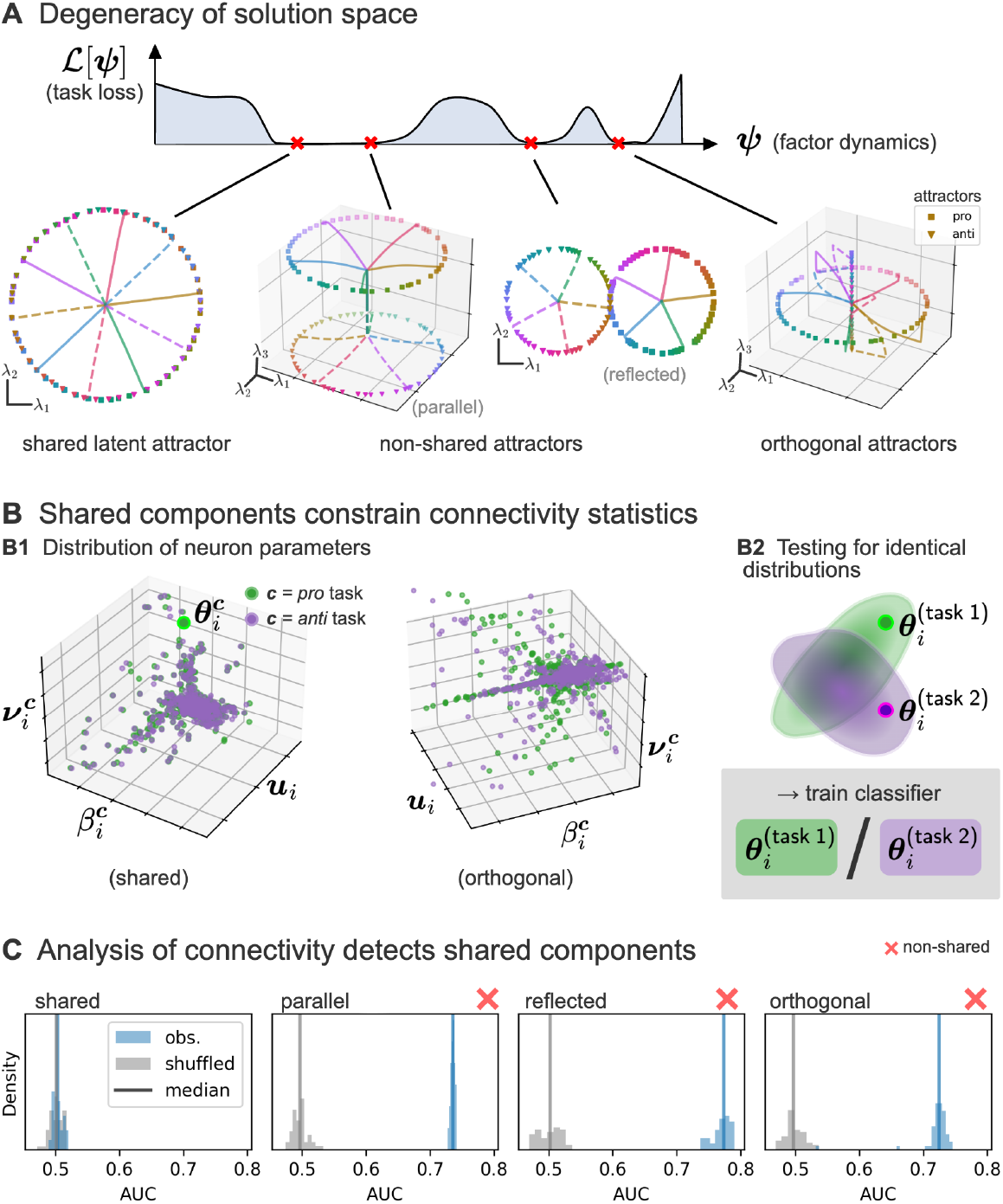
Shared components constrain network connectivity statistics. **A.** Many different solutions based on ring attractors in the latent dynamics exist for the memory pro/anti task, corresponding to minima in the sketched loss landscape. Instead of using a shared ring attractor, the network could use one ring attractor for each task which are separated along a third axis, reflected within the same subspace, or in orthogonal subspaces. **B**. Each neuron can be characterized by task-dependent effective parameters, which depend on the connectivity and the task input. Solutions using shared components imply identical statistics of the task-dependent effective neuron parameters (B1). This can be used to dissociate solutions with and without shared components by training a classifier to distinguish the distributions (B2). **C**. If the network uses shared components, the classifier cannot distinguish the statistics of the task-dependent effective neuron parameters above chance level. Histograms: distribution of classifier AUCs (blue: observed, gray: control with shuffled labels). Solid lines: medians.

To this end, we consider the flow-field 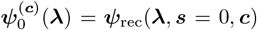 controlling the autonomous latent dynamics for a given task input ***c***, given from Eq. (3) by

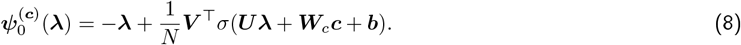

In general, the autonomous dynamics 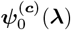 might depend on the task input ***c***. The fact that two tasks, with respective task inputs ***c*** and ***c***^′^, are solved using shared autonomous dynamics is formally expressed by the equality of the flow-fields 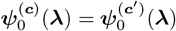. This equality imposes a subtle statistical structure in the network connectivity. To reveal this structure, we rewrite Eq. (8) as an empirical average over the neurons in the network [52–55],

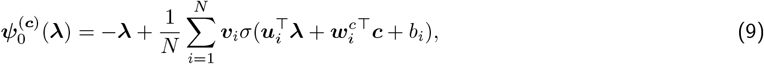

where the vectors ***u***_*i*_, ***v***_*i*_, 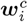 denote the *i*-th row of the matrices ***U***, ***V***, ***W***_*c*_ respectively, and *b*_*i*_ is neuron *i*’s bias. Formally, 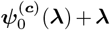 corresponds to the output of a single-hidden-layer feedforward network [56], where ***u***_*i*_, ***v***_*i*_ are input and output weights, and 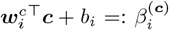 can be interpreted as an *effective* bias that is specific to task ***c***. Observe here that the task dependence enters only through the effective bias. The fact that two tasks ***c, c***^′^ share the same autonomous dynamics constrains the joint statistics of the input and output weights ***u***_*i*_, ***v***_*i*_ and effective biases 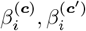. More precisely, for a given task ***c***, we define for each neuron *i* a task-dependent *effective output weight* 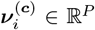, so that each neuron is characterized by a point 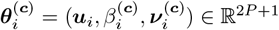 (Fig. 3B). We show that if two tasks ***c, c***^′^ use shared autonomous dynamics, then the point-clouds 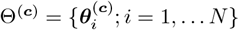 and 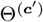 are statistically indistinguishable (Methods).

To illustrate this result, we designed three additional solutions to the pro/anti memory tasks presented above, which all use distinct ring attractors for the ‘pro’ and the ‘anti’ task (Fig. 3A). These distinct ring attractors are either shifted along a third dimension, reflected within the same plane, or in orthogonal subspaces. To test whether the point-clouds Θ^(***c***)^ are similar across the two tasks, with ***c*** corresponding either to the ‘pro’ task input ***c***_+_, or the ‘anti’ task input ***c***_−_, we trained classifiers to predict if a given point ***θ***_*i*_ was sampled from Θ^(***c***+)^ or Θ^(***c****−*)^ (Fig. 3B; Methods). As predicted by the theory, above chance classification, and thus statistical separation, is only possible for the RNNs using distinct attractors, as a hallmark that autonomous latent dynamics are significantly different between the two tasks (Fig. 3C).

### D. Hallmarks of shared attractors in representational geometry

We have shown that shared latent components constrain the connectivity statistics, although they are not immediately visible at the level of neural activity. However, synaptic connectivity is typically not accessible in behaving animals; thus, our method for identifying shared components by analyzing connectivity statistics is inadequate for an experimental setting. In contrast, neural activity can now be recorded at large scale across multiple tasks or contexts. Can shared latent components be detected from neural activity? We show that an analysis of neural activity during specific perturbation experiments provides a necessary condition to identify computation via shared attractors.

In these perturbation experiments, the task input is switched during a memory period: for a given pair of tasks with corresponding inputs ***c, c***^′^, the network is initially in an attractor state for task ***c***^′^, denoted 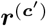. We then switch the task input from ***c***^′^ to ***c***, and let the network relax to a new attractor state, denoted 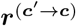 (Fig. 4A1). Finally, we compare this “perturbed” attractor state with another unperturbed attractor state ***r***^(***c***)^ for task ***c***. (Fig. 4A2; note that the task ***c*** of the unperturbed setting has to be the same as the final task in the perturbed setting ***c***^′^ → ***c***). We construct the corresponding representational similarity matrix (RSM) 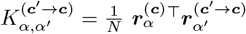, where *α* and *α*^′^ index different initial attractor states for each task, for example different points on the ring attractors of the ‘pro’ and ‘anti’ tasks. Intuitively, this matrix captures changes in the representational geometry [58] due to the perturbation of the task input from ***c***^′^ to ***c***. Reverting this procedure, we also determine the RSM 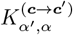 for the opposite perturbation from ***c*** to ***c***^′^.

**Figure 4.**
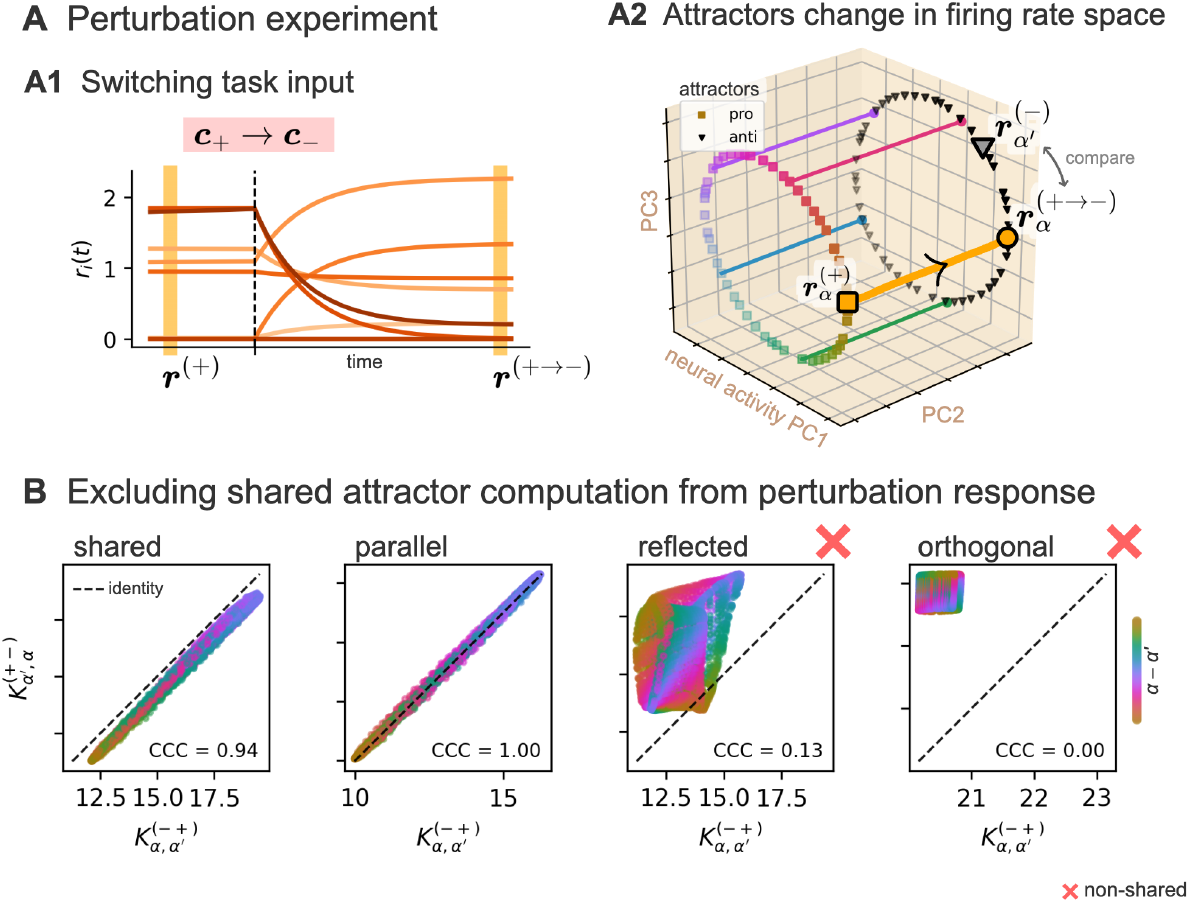
Perturbations of task input extract hallmarks of shared attractors in representational geometry. **A.** Starting from an attractor state ***r***^(+)^ for the ‘pro’ task, the task input is switched (vertical dashed line) from ***c***_+_ (‘pro’ task) to ***c***_−_ (‘anti’ task). This leads to firing rates dynamics towards a new “perturbed” attractor state ***r***^(+→−)^ (A1). Comparing ***r***^(+→−)^ with ‘anti’ task attractors ***r***^(−)^ across pairs of stimuli *α*^′^, *α* (A2) leads to the representational similarity matrix 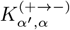. Colored solid lines: population activity trajectory after the task input switch ***c***_+_ → ***c***_−_ for five different initial attractor states 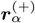. **B**. For shared attractors, the matrix entries 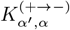 and 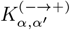 are identical. This symmetry allows to test for shared attractors across the four solutions from Fig. 3 (colored dots; color according to *α* − *α*^′^). For the “reflected” and “orthogonal” solution, the hypothesis of shared latent attractors can be excluded because of large deviations from the identity line. For the “parallel” solution the hypothesis can not be excluded, as a consequence of an additional symmetry of the solution. CCC: Lin’s Concordance Correlation Coefficient [57] between entries of the matrices 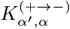 and 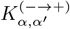.

For a shared attractor, the latent state is by definition not affected by the switch of the task input. In combination with the constraints imposed by shared latent dynamics on the connectivity statistics, we show that this implies a symmetry of the RSMs (Methods): 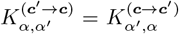. Importantly, this is only a necessary condition for a shared attractor; this symmetry could also arise from additional symmetries in the latent dynamics.

For a stylized example, we turn to the setup investigated in the previous section where the memory pro/anti task is solved with four different solutions (Fig. 3A). For the perturbation experiments, we switch the task input during the memory period; for instance, from the ‘pro’ task ***c***_+_ to the ‘anti’ task ***c***_−_. In all four solutions, the network stores the stimulus angle during the memory period using a ring attractor. After the perturbation and a sufficient relaxation period, the neural activity settles again in a state on a ring attractor. Even for the solution using a shared latent attractor, neural activity is affected by the perturbation, since shared attractors in latent space do not imply shared neural activity (Fig. 4A). However, changes in the representational geometry obey the symmetry 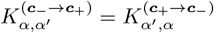 (Fig. 4B). We see explicitly that this symmetry is only a necessary condition: it is also exhibited by the solution with parallel ring attractors (Fig. 4B), due to its internal symmetries. This is, however, a peculiar example; in more generic cases we expect the symmetry to be violated for solutions relying on distinct attractors.

Going beyond examples with controlled solutions, we trained networks end-to-end on the same tasks, enforcing the rank of the recurrent connectivity matrix (Methods). Due to the training procedure, there is no explicit control over the solution found by the network. In particular, it is not clear whether the network utilizes shared latent attractors. Using either connectivity statistics or representational geometry reveals that occurrence of shared attractors depends on the hyper-parameters, in this case the rank (Fig. 5). This shows that either method can be used to identify shared attractors when the ground-truth solution is unknown.

**Figure 5.**
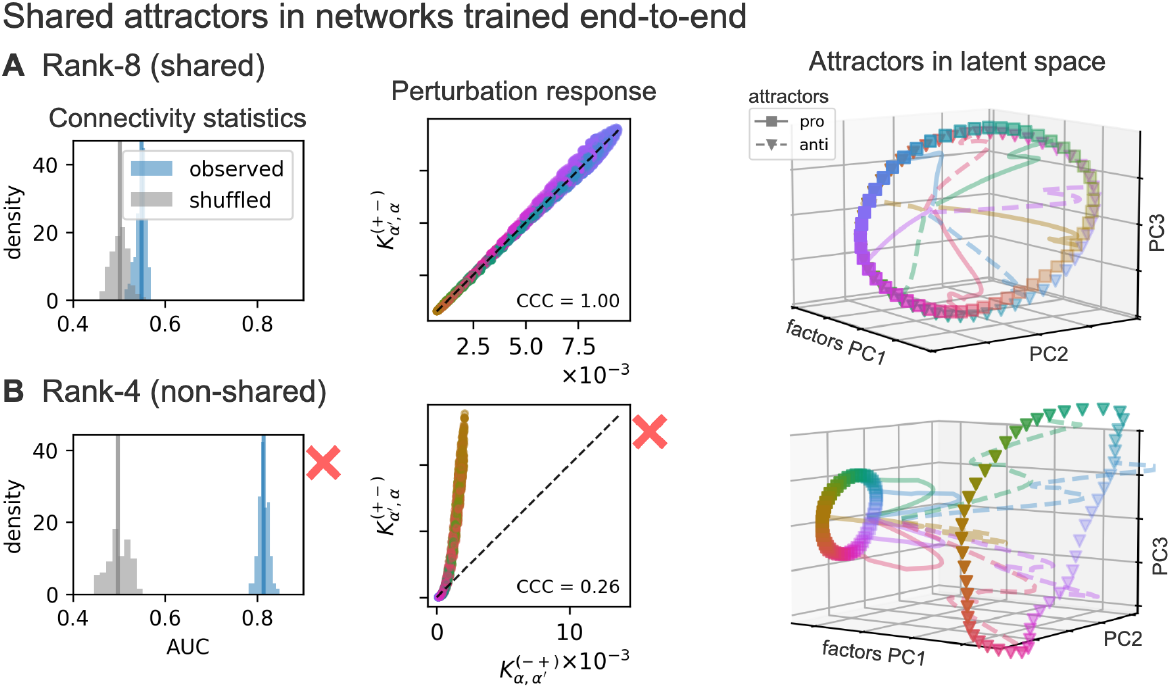
Identifying shared attractors in low-rank networks trained end-to-end. Low-rank networks were trained end-to-end on the pro/anti memory tasks (Methods). Analysis reveals whether networks use shared attractors (A, rank-8) or not (B, rank-4). Shared attractors are identified using either connectivity statistics (Fig. 3) or perturbation response (Fig. 4), and visualized with low-dimensional projections (PCA of factors ***λ*** during memory period).

### E. Loci of task-dependent computation

A central question about compositional computation is: where does the task-dependence enter the computation? Our framework identifies two potential loci. In a network using shared autonomous dynamics, the task input ***c*** can modulate how the stimulus drives the recurrent dynamics (via ***ψ***_in_, Eq. (5)), or how the shared factors are read out (via ***ψ***_out_, Eq. (4)). We call these two mechanisms *input-* and *output-modulation*. Task-dependence has to be resolved by one of the two, or a combination of both. For example, in the memory-pro/anti example above, we considered a solution via input-modulation: the shared ring attractor was used to store either the stimulus angle or its opposite, depending on the task input (Fig. 1C), while the readout was fixed across tasks. But this pair of tasks also admits a solution via output-modulation, where the input-driven dynamics (***ψ***_in_) are the same across the two tasks, but the readout is “flipped” for the anti-task. Both input- and output-modulated solutions rely on shared latent components. Importantly, the two potential implementations can be dissociated using perturbations of the task input: Switching the task input during the memory period changes the network’s output for the output-modulated, but not the input-modulated solution.

However, compositionality often relies on the combination of several independent contextual factors, which each affect the computation in a distinct way. To understand how the network can use shared components in this case, we move beyond the binary pro/anti task and turned to a family of tasks with an explicit compositional structure. Concretely, we consider a family of categorization tasks inspired by Tafazoli et al. [16]. In this study, macaque monkeys were trained to classify stimuli based on either their shape or their color, and had to report their decision using saccades along one of two orthogonal response directions. We modeled these tasks by representing the two stimulus features, color and shape, as two distinct input channels *s*_1_ and *s*_2_, and the two response directions as two output channels, *y*_1_ and *y*_2_. Each task in this family is therefore characterized by the combination of the relevant stimulus feature, *s*_1_ or *s*_2_ (indexed by a variable ***c***_in_), and the required response direction, *y*_1_ or *y*_2_ (indexed by ***c***_out_) (Fig. 6A). In contrast to the experimental setup, the stimulus presentation is followed in our model by a memory period and the network should respond once a fixation signal is switched off.

**Figure 6.**
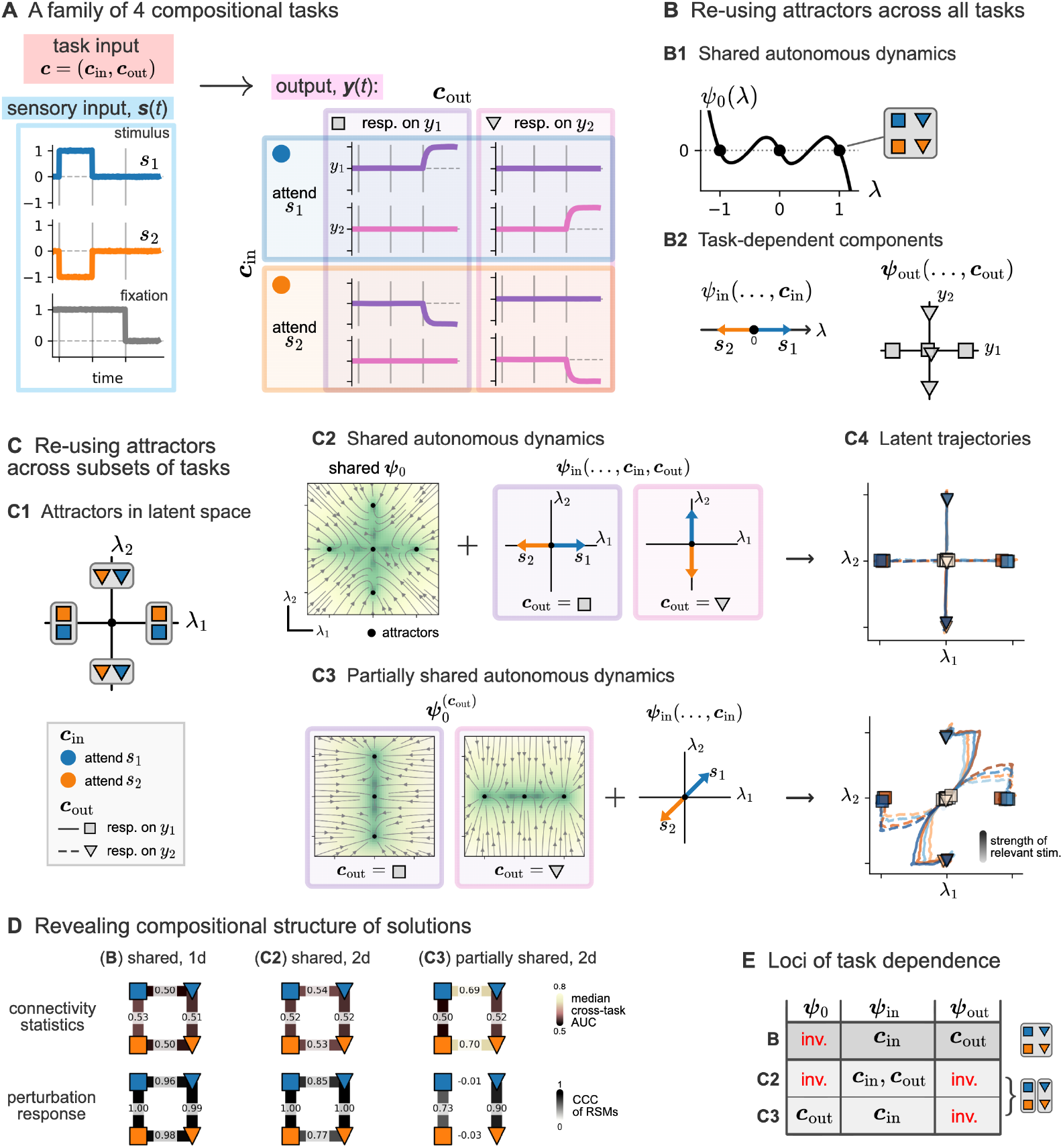
Combining shared and task-dependent components. **A.** A family of four categorization tasks where ***c***_in_ selects the attended input channel and ***c***_out_ the response channel reporting the categorization result. Each task is characterized by ***c***_in_ (blue or orange) and ***c***_out_ (square or triangular markers). **B**. Solution with attractors re-used across all tasks. (B1) Shared one-dimensional latent autonomous dynamics. Black dots: attractors, used to store the classification result. (B2) ***c***_in_-dependence is resolved by input-driven dynamics ***ψ***_in_; ***c***_out_-dependence is resolved by readout ***ψ***_out_. Left: colored arrows represent the value of ***ψ***_in_ at *λ* = 0 for the stimulus of panel (A) and the two values of ***c***_in_ (blue or orange). Right: markers represent ***ψ***_out_ evaluated at the three attractors of panel (B1) for the two values of ***c***_out_ (squares or triangles). **C**. Solution with attractors re-used across subsets of tasks. (C1) Latent attractors used for each task. Output-dependence is resolved by recurrent dynamics: attractors are re-used only across tasks with the same ***c***_out_. Autonomous dynamics can be shared across tasks (C2), or depend on ***c***_out_ (C3). In the former case the input-driven dynamics must resolve the full task-dependence; in the latter they only have to resolve ***c***_in_. Colored arrows as in (B2). (C4) Both solutions lead to distinct latent dynamics. Solid and dotted lines: trajectories; markers: fixed points. **D**. Identification of shared components across tasks for each solution. Markers represent tasks. Pairs of tasks (edges) are compared using connectivity statistics and measuring median cross-task classifier AUC (see Fig. 3); or using task input perturbations and measuring Lin’s CCC of representational similarity matrices (RSMs; see Fig. 4). High AUCs and low CCCs indicate non-shared autonomous dynamics. Non-shared components are successfully identified (light edges). **E**. Overview of the loci of task-dependence. For each solution (rows), entries summarize which component of the latent dynamics (***ψ***_0_, ***ψ***_in_ or ***ψ***_out_, columns) resolves each task-dependence, ***c***_in_ and ***c***_out_. “inv.”: task-invariant.

Similar to the memory pro/anti task, there are different solutions based on shared latent components. In a minimal solution, the categorization choice is encoded using only two attractors, which are re-used across all task conditions (Fig. 6B1). This solution combines input- and output-modulation: the input flow-field selects the relevant stimulus feature according to ***c***_in_, and the readout flow-field directs the response to the required direction according to ***c***_out_ (Fig. 6B2), while the autonomous dynamics are fully invariant to the task input.

However, the step from a single task condition to multiple, compositional ones opens up a set of multiple qualitatively different solutions: instead of using a shared component across all tasks, specific components could be shared across subsets of tasks. As an example, we constructed solutions that use orthogonal latent attractors across different response directions; in this way, the choice of the response direction is delegated to the recurrent dynamics while the readout ***ψ***_out_ can be shared across all tasks. In these solutions, attractors are re-used across conditions for the relevant stimulus feature (***c***_in_), but not for the selected response direction (***c***_out_). This is possible in a two-dimensional latent space, in which each combination of categorization choice and response direction corresponds to one of four different attractors (Fig. 6C1). The autonomous dynamics can be shared across all tasks, with four coexisting attractors (Fig. 6C2). Alternatively, the autonomous dynamics could only contain the two attractors corresponding to the selected response direction ***c***_out_, while the attractors for the non-selected direction are absent (Fig. 6C3). In the latter case, the autonomous dynamics are only *partially shared* across tasks; concretely, they are invariant with respect to ***c***_in_ but depend on ***c***_out_. In full analogy to the fully shared components considered in the previous sections, these partially shared components lead to testable hallmarks in the connectivity and representational geometry (Fig. 6D). Furthermore, they can be distinguished using perturbations of the task input during the memory period: changing ***c***_out_ during a memory period changes the network output for the ‘partially shared’ solution, but not for the ‘fully shared’ one.

Each of the three solutions presented above is distinguished by the locus where the selection of the output direction enters the computation (Fig. 6E). In the one-dimensional solution, this is performed at the level of the readout ***ψ***_out_; in the ‘fully shared’ solution, this is done via the input flow-field ***ψ***_in_; and in the ‘partially shared’ solution, via the autonomous flow-field 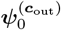. For all three solutions, input feature selection is performed by ***ψ***_in_. Building on this classification of solutions by the locus of task-dependence, we systematically constructed a family of nine qualitatively different solutions, each uniquely characterized by the structure of attractor re-use and the specific combination of the loci (***ψ***_0_, ***ψ***_in_, or ***ψ***_out_) associated with each task condition (***c***_in_ and ***c***_out_) (Methods and Fig. S1).

### F. Compositional computation via shared latent dynamics

In this section, we show explicitly how shared and task-dependent components can be composed to solve a more elaborate family of tasks, going beyond the stylized examples employed so far. Here, we use fully shared autonomous dynamics to demonstrate how they can be combined with task-specific components at different loci to enable multi-tasking.

Concretely, we consider a family comprising eight of the tasks used by Driscoll et al. [15]: delay-pro, memory-pro, react-pro, delay-anti, memory-anti, react-anti, match-to-sample, and anti-match-to-sample (Fig. 7A1; see the Methods for detailed task descriptions). The delay and react tasks are similar to the memory tasks used above, except that there is no memory period: after its onset, the stimulus is continuously present. For the memory and delay tasks, the output should occur when the fixation input switches off; for the react task it should occur directly at stimulus presentation. The match-to-sample tasks are structurally different: in these tasks the network receives a second stimulus after a memory period, and should report the second stimulus if it matches the first one without a delay.

**Figure 7.**
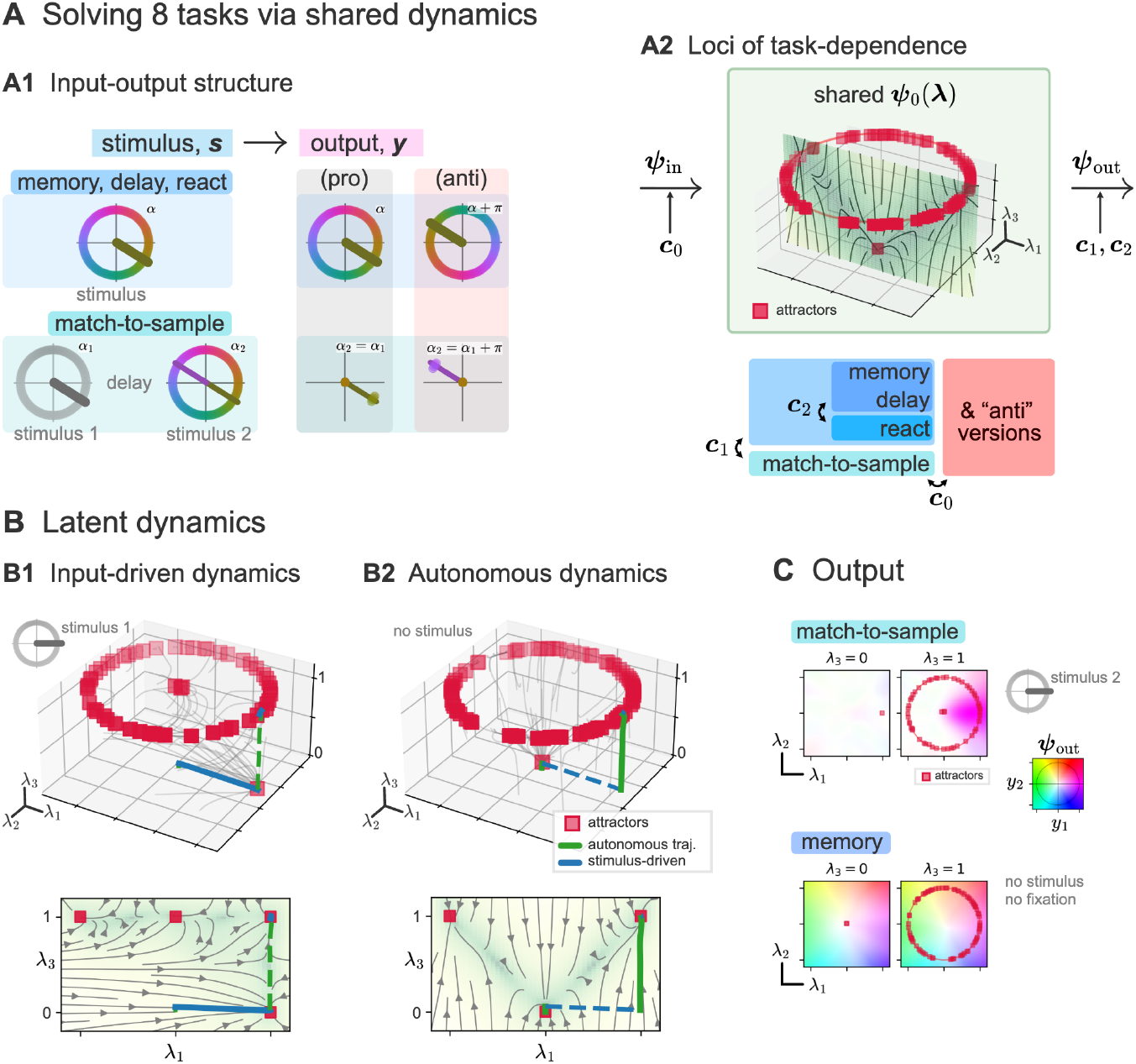
Compositional computation across eight tasks via shared latent dynamics. **A1.** Input-output structure for the eight tasks. Memory, delay, and react require reporting a stimulus angle *α* (pro) or its opposite *α* + *π* (anti); they differ in stimulus presentation timing and output gating (Methods). Match-to-sample tasks require comparing a second stimulus angle *α*_2_ to a memorized first one *α*_1_ and reporting a match (pro: *α*_2_ = *α*_1_, anti: *α*_2_ = *α*_1_ + *π*). **A2**. The eight tasks are solved using a shared autonomous flow field ***ψ***_0_, comprising a fixed point and a ring attractor in a three-dimensional latent space. Task-dependence is resolved by the input flow field ***ψ***_in_ (modulated by ***c***_0_, encoding pro/anti) and the readout ***ψ***_out_ (modulated by ***c***_1_, ***c***_2_, encoding task class). Flow-field in green shows dynamics in the (*λ*_2_ = 0) plane. **B**. Input-driven and autonomous dynamics. **B1**. An incoming stimulus drives the network towards the corresponding location in the *λ*_3_ = 0 plane (blue trajectory). Bottom: flow-field in the *λ*_2_ = 0 plane. **B2**. During the memory period, autonomous dynamics drive the network to the ring attractor in the *λ*_3_ = 1 plane, retaining the stored angle (green trajectory). **C**. Readout ***ψ***_out_, color-coded (white: zero output). For the match-to-sample task (top), output is produced only in the *λ*_3_ = 1 plane, where the location on the ring matches the second stimulus. For the memory task (bottom), the readout matches the latents (*λ*_1_, *λ*_2_).

Our network implementation solves this family of tasks using a three-dimensional autonomous flow-field that contains a stable fixed point at the origin and a ring attractor that is shifted in the z-direction (Fig. 7A2). At the beginning of the task, the network is in the fixed point at the origin. An incoming stimulus and the subsequent autonomous dynamics (Fig. 7B) drive the network to the corresponding location on the ring attractor, or its opposite for anti tasks. Importantly, the third factor, *λ*_3_, acts as a gate: the stimulus affects the dynamics only in the *λ*_3_ = 0 plane, but not on the ring attractor in the *λ*_3_ = 1 plane. This gating allows the network to solve the match-to-sample tasks: the first stimulus, which is stored on the ring attractor, is not affected by the incoming second stimulus, such that both stimuli can be compared by the readout.

The solution combines multiple loci to resolve task-dependence (Fig. 7A2). The pro vs. anti context is resolved by the input flow-field: in the anti context, the input-driven dynamics implement a rotation by *π* for all tasks. The readout flow-field resolves the four classes of tasks: for the match-to-sample tasks, the readout compares the incoming second stimulus to the stored first stimulus and reports it if there is a match (Fig. 7C). Put differently, the readout is non-zero only if the network is on the ring attractor (*λ*_3_ = 1) and the incoming stimulus matches the stored angle. For the memory, delay, and react tasks, the readout simply reports the value of the two first latents, encoding the stimulus angle, or its opposite for anti tasks. In addition, the readout is gated by the fixation signal for the memory and delay tasks.

This example shows how a recurrent network can solve a range of different tasks with a common structure using shared autonomous latent dynamics, with all task-dependence delegated to input and readout flow-fields. In this case, the shared structure is that all tasks rely on an angular variable, which is naturally implemented by a shared ring attractor.

## III. DISCUSSION

Building on empirical results in trained recurrent neural networks [12, 14, 15], we developed a theory of compositional computation via shared latent components in low-rank recurrent networks. In particular, we proposed that the notion of “shared dynamical motifs”, introduced by Driscoll et al. [15], can be formalized in terms of autonomous latent dynamics that are shared and re-used across tasks involving similar computations. We showed that these shared latent components are consistent with task-dependent neural activity, which is ubiquitously observed in neural data [17, 24, 25, 30, 46–51]. Furthermore, we used our theory to derive observable consequences of computation via shared latent components. First, we showed that the connectivity statistics contain signatures that allows to dissociate shared from non-shared solutions. Second, we utilized perturbations of the task input to reveal hallmarks in the representational geometry imposed by shared attractors. Finally, we explicitly constructed solutions for compositional tasks to demonstrate that shared latent components enable flexible and dynamic compositional computation. Based on these solutions, we delineated specific mechanisms through which the shared components can be utilized: via input- or output-modulation, and via partial invariances.

Our work highlights the degeneracy of solutions in recurrent neural networks [14, 59–61]. The low-rank assumption for the network connectivity poses a strong constraint on the space of possible networks. Nonetheless, many low-rank solutions are possible for each task we considered; we explicitly constructed some of these solutions. For large enough networks, the universal approximation property of feedforward networks [62–64] together with the mapping from low-rank recurrent to feedforward networks [56] implies that arbitrary solutions with the dimensionality given by the rank of the connectivity can be implemented.

The degeneracy of the solution space raises the question of how these solutions can be distinguished. Based on our theory, we showed that targeted perturbations of the task input provide a powerful tool to dissociate solutions. Importantly, this approach relies only on experimentally accessible quantities: the task signal and the neural activity. Hence, our theory suggests new experiments where targeted perturbation of the task- or context-signal are used to gain insight into the mechanisms employed by the network. Note that using perturbation of the task signal is different from using perturbations of the stimulus inputs [50], neural activity [65], or connectivity [44] to dissociate solutions.

A generic problem for compositional tasks is how a network can discover and exploit the tasks’ compositional structure during training. This problem is tightly related to the encoding of the task identity [13]: if the input directly encodes the compositional structure, it can be exploited by the network to enable compositional generalization; the opposite extreme is a one-hot coding which does not contain any information about the task structure. For our theory, this distinction is not relevant. In both cases, the context input reduces to a task-dependent effective bias. From this perspective, structured compositional inputs and shared components form distinct, complementary mechanisms which facilitate compositional computation. Importantly, we assume for all our examples that the correct task is encoded in the input, akin to a task set [66, 67], circumventing the challenging question how the correct task can be inferred [68].

Conceptually, we leverage the low-rank assumption and a strict mathematical definition of shared latent components to derive observable consequences. While end-to-end trained neural networks do not necessarily obey the low-rank assumption, training often leads to learned low-rank components [35], such that the networks might rely on qualitatively similar dynamics. A direct consequence of the low-rank assumption is that the neural activity is constrained to low-dimensional manifolds [69, 70] which are nonlinearly embedded in the firing rate space (cf. Eq. (7)). Due to the nonlinear mapping, the resulting manifolds can have a high linear dimensionality in firing rate space [71], as observed in neural activity [72, 73]. An alternative theoretical approach to multi-task networks with high-dimensional dynamics does not use task-shared dynamical components [74].

A central component of our framework is the mean-field description of single-hidden-layer networks in terms of a probability distribution [52–55]. In the context of low-rank networks, this distribution is typically assumed to be Gaussian [27] or a mixture of Gaussians [31, 37, 45]; recent studies went beyond Gaussianity to link connectivity and computation in the case of more general connectivity structures [43, 75]. Going beyond the (mixture of) Gaussian assumption increases the expressivity of the network and links to universal approximation, which we expect to be of broad interest in theoretical neuroscience.

Existing work on multitasking recurrent networks relies predominantly on end-to-end training. Here, we took an orthogonal approach: we designed flow-fields that solve the task and trained feedforward networks to implement them, exploiting the link between low-rank recurrent networks and feedforward networks [56]. This allowed us to have precise control over the solution implemented by the network; in contrast, end-to-end training relies on a variety of indirect methods, for example hyperparameter tuning, to chart the solution space [14, 15, 59–61, 76]. On a more abstract level, controlling the flow-field allowed us to strike a favorable balance that retains mechanistic interpretability while allowing for complex solutions to multi-task problems.

## Acknowledgements

We thank Wulfram Gerstner for exceptional guidance and mentoring, Manuel Beiran for helpful feedback on the project, Flavio Martinelli for valuable discussions on identifiability, and the whole team of the Laboratory of Computational Neuroscience for insightful discussions. We gratefully acknowledge the publicly available data by Valerio Mante and Laura Driscoll that is used in Fig. 2A. This work was funded by the Swiss National Science Foundation grants CRSII5 198612, 200020-207426 and 200021-236436. AvM was also supported by the Ministry of Culture and Science of North Rhine-Westphalia (NRW Rückkehrprogramm).

## IV. METHODS

### A. Neural network training

#### 1. Controlling solution space degeneracy

End-to-end training of recurrent neural networks uses (stochastic) gradient descent to minimize a loss function ℒ(Θ) on the space of the network parameters Θ, for example a mean-squared-error loss that penalizes deviations from the desired network output [15]. Typically, there are multiple, qualitatively different solutions to the task [14, 59–61, 76]. Put differently, the solution space exhibits a degeneracy. Which of these solutions is found depends on the implicit bias imposed by, among other factors, hyperparameters, initialization, network architecture, and learning algorithm.

For low-rank networks, this solution space degeneracy can be controlled. The central observation is that the network output depends on the parameters only through the flow fields, see (3) and (4). Hence, in this case one can consider the loss as a functional of the flow fields ℒ [***ψ***], where we denote ***ψ*** = {***ψ***_rec_, ***ψ***_out_} for simplicity. To control the solution space degeneracy, we introduce a target flow field ***ψ***_∗_ corresponding to a specific solution to the task, ℒ [***ψ***_∗_] ≈0. Controlling the properties of the target flow field allows us to control the properties of the solution. The target flow field, in turn, is imposed using an auxiliary mean-squared-error loss,

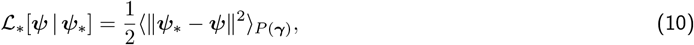

where *P* (***γ***) denotes the data distribution comprising the latent variables as well as all input variables. Adjusting the data distribution allows to control the domain within which the flow field of the network matches the target flow field; in regions of low probability *P* (***γ***) the flow field can deviate from the target flow field.

To train the flow field, we use online stochastic gradient descent (SGD) with L2 regularization and Gaussian noise. New data samples are generated from *P* (***γ***) in each SGD step. For large network sizes *N*, this training procedure is described by a convex optimization problem in the space of probability measures of the single-neuron parameters *ρ*(***θ***), which implies that it is guaranteed to converge [52–55]. Intuitively, the convexity arises because the flow field ***ψ*** can be written as a linear functional of *ρ*, such that the auxiliary mean-squared error loss ℒ_∗_[***ψ*** |***ψ***_∗_] is quadratic in *ρ* [52].

Conceptually, our training approach is fundamentally different from end-to-end training: leveraging the correspondence between low-rank networks and the low-dimensional latent dynamics described by ***ψ***, we specify a solution to the task ***ψ***_∗_. To train the flow field, we need to train the corresponding single-hidden-layer network, (3) and (4), on the auxiliary MSE loss (10). Note that this is computationally cheap compared to end-to-end training with backpropagation through time.

#### 2. Flow field training

For each task solution used in the Results, we designed a specific flow-field (***ψ***_rec∗_, ***ψ***_out∗_), see section “Task solutions”. To obtain low-rank networks implementing the dynamics of Eqs. (3)-(4) with given (***ψ***_rec∗_, ***ψ***_out∗_), we trained MLPs on the auxiliary loss of Eq. (10), with the target ***ψ***_∗_ defined as ***ψ***_∗_(***λ, s, c***) = (***ψ***_rec∗_(***λ, s, c***) + ***λ, ψ***_out∗_(***λ, s, c***)).

All MLPs were trained using online SGD. At each step, a new sample was drawn independently from the task-specific training data distribution *P* (***λ, s, c***), see section “Task solutions”. L2 regularization with coefficient *λ*_reg_ was applied to all weights. Weight diffusion was implemented as additive Gaussian noise with standard deviation *σ*_diff_, applied independently to each weight at every gradient step. Gaussian noise with standard deviation *σ*_in_ was further added to the inputs ***s, c*** to improve robustness. Numerical parameters were set to 1.5 × 10^7^ gradient steps (except ~40 × 10^7^ for the 8-tasks flow field of Fig. 7), step size *η* = 5 ×10^−3^, *λ*_reg_ = 10^−6^, *σ*_diff_ = 10^−6^, *σ*_in_ = 10^−6^. Number of neurons was *N* = 1024 (except *N* = 2048 for Fig. 7).

#### 3. End-to-end training

We trained two low-rank networks end-to-end on the pro/anti memory tasks, with ranks *P* = 8 and *P* = 4 (Fig. 5). Empirically, low-rank networks initialized at random did not reliably converge; we therefore used a two-stage procedure. First, a full-rank network of *N* = 256 neurons was trained end-to-end on the two tasks. Second, its recurrent connectivity was truncated to rank *P* by retaining the *P* leading components of its singular value decomposition, 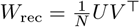; this low-rank approximation was used to initialize a low-rank network, which was then fine-tuned end-to-end on the same tasks.

Both stages used backpropagation through time with the Adam optimizer and a mean-squared-error loss on the network output. At each step, a fresh batch of 64 trials was drawn from the task distribution. The discretized dynamics used a step size d*t/τ* = 0.2. L2 regularization with coefficient *λ*_reg_ = 10^−6^ was applied to the weights, and an additional penalty with coefficient *λ*_act_ = 10^−6^ to the firing rates. Gaussian noise was injected into the inputs (*σ*_in_ = 0.1) and into the recurrent currents (*σ*_rec_ = 0.05) at every step to improve robustness. Both full-rank pretraining and low-rank fine-tuning used a learning rate *η* = 3 × 10^−4^ and 2 × 10^5^ steps for the rank-8 network, and *η* = 10^−3^ for 10^5^ steps for the rank-4 network.

### B. Shared components impose constraints on the empirical measure of parameters

We start from the expression of Eq. (9) for the autonomous dynamics. Up to the leak term (−***λ***), this expression can be interpreted as a multi-layer perceptron (MLP) with a single hidden layer, where the term 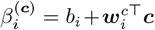 is equivalent to a task-dependent bias. Thus, the problem can be mapped to an identifiability problem for MLPs: How does the input-output function of an MLP constrain its parameters?

A MLP with *N* hidden neurons is defined by a nonlinearity *σ*: ℝ → ℝ, and a set of parameters Θ = {(***u***_*i*_, ***v***_*i*_, *β*_*i*_), *i* = 1, …, *N*}, where ***u***_*i*_ denotes neuron *i*’s input weights, ***v***_*i*_ its output weights, and *β*_*i*_ its bias, parameterizing the function

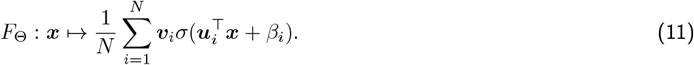

MLPs can *approximate* arbitrary functions if the number *N* of neurons is large enough due to the Universal Approximation Theorem [62–64]. A distinct question is that of *identifiability*: can two MLPs with different parameters implement the same function? In other words, are network parameters uniquely determined by the function they implement? Identifiability results have been derived in various settings [77–79], including networks with more general architectures [80, 81].

#### 1. Symmetries, irreducibility, and empirical measure

Importantly, Eq. (11) enjoys general symmetries beyond the well-known permutation symmetry of neuron labels, that must be taken into account when considering identifiability [77, 79, 82]:

1. *Duplicate neurons*: if several neurons *i*_1_, *i*_2_, …, *i*_*m*_ have the same input weights and biases 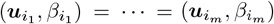, the function *F*_Θ_ does not depend on their individual output weights 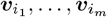 but only on their sum 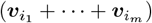.
2. *Zero neurons*: if a neuron’s output weight is null, ***v***_*i*_ = 0, the function *F*_Θ_ does not depend on its other parameters ***u***_*i*_ and *β*_*i*_.

Therefore, identifiability results are typically expressed for “irreducible” networks containing neither zero nor duplicate neurons.

Alternatively, Eq. (11) can be expressed in a way that naturally encompasses the permutation, duplicate-neurons, and zero-neurons symmetries. To this end, we start by writing it in terms of the empirical measure (EM) of the single-neuron parameters

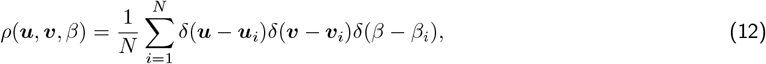

where *δ*(*x*) denotes a Dirac delta, as

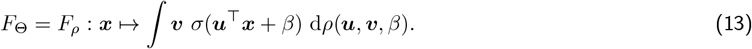

By decomposing the EM as the product of its (***u***, *β*)-marginal *µ*(***u***, *β*) with the corresponding conditional *ρ*(***v***|***u***, *β*), one can write

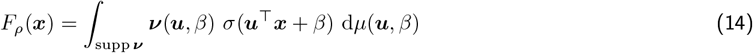

where ***ν***(***u***, *β*) := ***v*** d*ρ*(***v*** |***u***, *β*) denotes the conditional mean of the output weights ***v***, and supp ***ν*** its support. The function *F*_*ρ*_ neither depends on the conditional distribution *ρ*(***v*** |***u***, *β*) beyond its mean (see also [83]), nor on the distribution *µ* outside the support of ***ν***: these invariances account, respectively, for the *duplicate-neurons* and *zero-neurons* symmetries. Therefore, the identifiability of irreducible MLPs is equivalent to the identifiability of the marginal *µ* and conditional mean ***ν*** for non-irreducible MLPs: if two MLPs with EMs *ρ* and *ρ*^′^ implement the same function *F*_*ρ*_ = *F*_*ρ*_*′*, then ***ν*** = ***ν***^′^ and *µ* = *µ*^′^ on supp ***ν***.

The properties of the activation function may give rise to additional symmetries. For example, softplus *σ*(*t*) = log(1 + *e*^*t*^) has the “even+linear” property *σ*(*t*) = *σ*(−*t*) + *t* [82]. Such activation-function-specific symmetries can in turn lead to *affine symmetries* [81], i.e., non-trivial combinations of neurons *I* with a constant output, 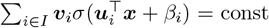 const. Crucially, Vlačić and Bölcskei [81] showed that irreducible MLPs are identifiable up to permutation if *σ* has no such affine symmetries; and conversely, affine symmetries are the only additional symmetries beyond the generic permutation, zero-neurons, and duplicate-neurons symmetries.

In summary, MLPs are identifiable up to specific symmetries. These symmetries are conveniently captured by the parameterization of Eq. (14), excluding possible activation-function-specific affine symmetries.

#### 2. Approximate conditional mean

To distinguish shared and non-shared latent dynamics, we test for statistical difference between the empirical samples Θ for two tasks. Since the order of the samples is not relevant for the test, this takes the permutation invariance into account. Furthermore, we estimate the conditional mean of the output weights ***ν***(***z***) to account for the duplicate-neurons symmetry. For simplicity, we do not consider the zero-neuron symmetry as well as additional activation-function-specific affine symmetries. Empirically, zero-neuron symmetries are absent, likely due to weight diffusion during training, and we expect that the involved affine symmetries occur only rarely.

We denote the input weights and the effective bias as 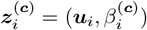, and drop the superscript ***c*** unless it is relevant. The empirical conditional mean of the output weights ***ν***(***z***) ≡ ∫ ***v*** d*ρ*(***v***| ***z***) reads

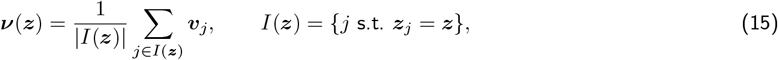

which is only defined for values of ***z*** such that there is at least one neuron with ***z***_*j*_ = ***z***, i.e., *I*(***z***) *>* 0. If there are no duplicates the conditional mean is simply ***ν***(***z***_*i*_) = ***v***_*i*_.

Empirically, we did not observe exact duplicates in ***z***-space. This may be linked to numerical imprecision as well as weight diffusion during training. We interpret this fact in a statistical perspective: redundancy in ***z***-space has to be understood in a smooth way, where neurons close to each other are *almost* duplicates. To account for this perspective, we smoothen the Dirac deltas of the EM in Eq. (12) using a Kernel Density Estimate (KDE) of *ρ* [84] to compute an approximate conditional mean: Let 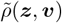 be a KDE of the EM *ρ*, of the form

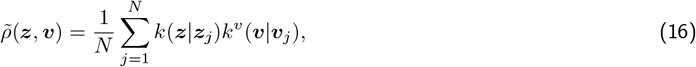

where *k*(·|***z***^∗^) is a kernel with mass 1 and mean ***z***^∗^ (e.g., a Gaussian), and similarly for *k*^*v*^(·|***v***^∗^). We compute an approximate conditional mean as

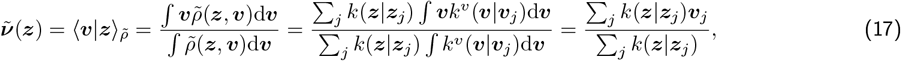

which yields an “effective output weight” for each neuron *i*,

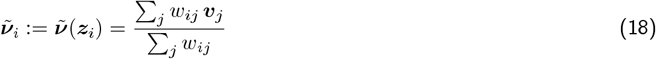

where *w*_*ij*_ = *k*(***z***_*i*_ | ***z***_*j*_). Note that the choice of the kernel *k*^*v*^(·| ***v***^∗^) is irrelevant here, provided that it has mass 1 and mean ***v***^∗^.

For a given task ***c***, let 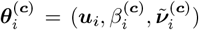 denote the single-neuron parameters that characterize the autonomous dynamics, and 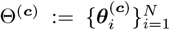 the corresponding point-cloud. If the point-clouds Θ^(***c***)^ and 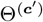 are statistically undistinguishable for two tasks ***c, c***^′^, the tasks share similar autonomous dynamics. Conversely, if the point-clouds are significantly different, either the (***u***, *β*^(***c***)^)-marginals, or the function 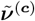, are different across tasks — which implies that autonomous dynamics are different across tasks due to the identifiability of MLPs expressed in terms of the (***u***, *β*)-marginal and the conditional mean.

For the KDE of *ρ*, we used Gaussian kernels of the form 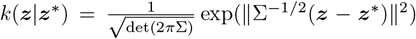 where 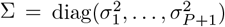 is a diagonal matrix with separate bandwidths *σ*_*µ*_ for each dimension *µ*. To select the maximal bandwidth, we used that the autonomous flow-field should not be significantly affected by replacing each ***v***_*i*_ with the corresponding 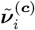. For instance, for the pro/anti networks with a shared ring attractor, this replacement should not destroy the slow ring manifold. Concretely, bandwidth selection was made according to the scale of the data: bandwidth *σ*_*µ*_ in dimension *µ* was set to 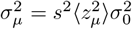, with 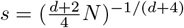 a proportionality factor that adjusts the bandwidth to the dimensionality *d* of ***z*** and the number *N* of neurons (following Silverman’s ‘rule of thumb’ for multivariate data [84]), 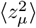 the mean square of samples of *z*_*µ*_, and 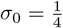 a fixed factor small enough to ensure preservation of flow fields.

#### 3. Testing for statistical difference between point-clouds

Given two point-clouds Θ^(***c***)^ and Θ^(***c***^*′*), we tested whether there was a significant difference between their underlying distribution by training nonlinear classifiers to predict from which point cloud (Θ^(***c***)^ or Θ^(***c***^*′*)) a given data point ***θ*** was sampled [85, 86].

Importantly, the (***u***)-marginals of the point-clouds are identical because the input weights {***u***_*i*_} are the same across tasks. This can lead to identity leak when training classifiers if the same ***u***_*i*_ is shared between training and test sets. To prevent such an identity leak, cross-validation folds were defined over neuron indices *i*: for each fold, both data points associated with index *i* (i.e. both 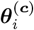 and 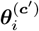, with label *y*_*i*_ = +1 or −1 respectively) were assigned together to either the training or test set. Classifiers were trained to predict labels from data. The AUC was computed by averaging over folds. Tests under the null hypothesis were performed by independently swapping labels for each neuron with probability 1*/*2, which preserves the marginal distributions while destroying the correspondence between class labels and parameter features.

To obtain the results reported in the figures, we trained nonlinear Support Vector Classifiers (SVCs) with an RBF kernel using scikit-learn [87] and evaluated their AUC using 2-fold cross-validation. A distribution of AUCs was obtained by running cross-validation over 50 different classifier initializations and random fold assignments. When reporting a single AUC (as in Fig. 6D), it corresponds to the median of this distribution.

### C. Testing for shared latent attractors via task input perturbations

Given a latent stationary state ***λ*** of the autonomous dynamics, the corresponding population activity of neuron *i* for task input ***c*** is given from Eq. (7) by

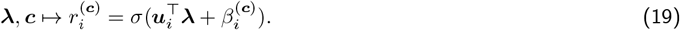

Let 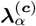 be an attractor of the autonomous latent dynamics for task input ***c*** and sensory input *α*, and 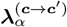 the new attractor reached after a switch of the task input from task input ***c*** to ***c***^′^. Denote 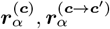 the corresponding population activity vectors. We define 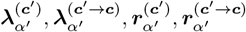 similarly, for the task switch from ***c***^′^ to ***c*** and the sensory input *α*^′^.

Define the kernel (also called representational similarity matrix) for two inputs *α, α*^′^,

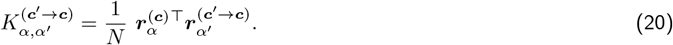

If ***c, c***^′^ share autonomous dynamics, then (i) 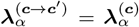, and (ii) the marginal distributions coincide, 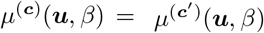 Therefore,

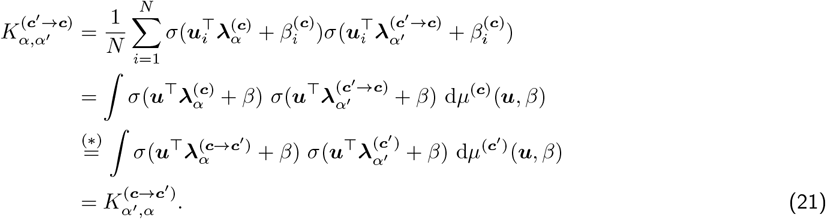

where we used (i) and (ii) in (∗). Thus, the symmetry 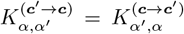 is a direct consequence of shared latent attractors. Conversely, a violation of this symmetry indicates an absence of shared attractors.

### D. Tasks

#### Memory pro / anti tasks

This family comprises two tasks: memory pro and memory anti. The input is four dimensional, *s* = (*s*_1_, *s*_2_, *f, c*) where (*s*_1_, *s*_2_) ∈ ℝ^2^ are the stimulus input, *f* ∈ {0, 1} the fixation input, and *c* ∈{−1, 1} encodes the pro (*c* = 1) or anti (*c* = −1) version of the tasks; the target output is two-dimensional, *y* = (*y*_1_, *y*_2_) ∈ ℝ^2^. There are four consecutive periods: context, stimulus, memory, and response (following Driscoll et al. [15]). The stimulus input is (*s*_1_, *s*_2_) = (cos *θ*_*s*_, sin *θ*_*s*_) with *θ*_*s*_ ∈ [0, 2*π*) during the stimulus period and zero otherwise, the fixation input is *f* = 0 during the response period and *f* = 1 in all other previous periods, and the context input is constant (*c* = ±1) during all periods. The target output (Fig. 1A) is (*y*_1_, *y*_2_) = (cos *θ*_*s*_, sin *θ*_*s*_) or (*y*_1_, *y*_2_) = (cos(*θ*_*s*_ + *π*), sin(*θ*_*s*_ + *π*)) for pro and anti tasks, respectively, during the response period and zero during other periods.

For simulations, periods ranged across 0.5 to 1.5*τ* for the context, 2 to 5*τ* for the stimulus, 5 to 10*τ* for the memory, and 5*τ* for the response.

#### Ring attractor family

This family comprises eight tasks: react, delay, memory, and react-to-sample, each in a pro and anti version. The input is six dimensional, *s* = (*s*_1_, *s*_2_, *f, c*_0_, *c*_1_, *c*_2_), where *c*_1_ ∈ {0, 1} encodes whether the match-to-sample (*c*_1_ = 1) or another task (*c*_1_ = 0) should be solved, *c*_2_ ∈ {0, 1} distinguishes react (*c*_2_ = 1) from delay and memory tasks (*c*_2_ = 0), and *c*_0_ encodes the pro/anti context (as *c* in the memory tasks above). The target output is two-dimensional, *y* = (*y*_1_, *y*_2_) ∈ ℝ^2^. There are two consecutive periods in the react tasks (context and stimulus), three periods in the delay tasks (context, stimulus, and response), and four periods in the memory and match-to-sample tasks (context, stimulus, memory, and response). All task inputs are constant during all periods; the fixation input is *f* = 1 except for tasks with a response period during which *f* = 0. The stimulus is (*s*_1_, *s*_2_) = (cos *θ*_*s*_, sin *θ*_*s*_) during stimulus (react, memory, and match-to-sample tasks) or stimulus and response (delay tasks) periods; for the match-to-sample tasks a second stimulus (*s*_1_, *s*_2_) = (cos *θ*_*m*_, sin *θ*_*m*_) is presented during the response period; and during all other periods the stimulus is zero.

Similar to the memory pro/anti tasks, for the react and delay tasks the target is (*y*_1_, *y*_2_) = (cos *θ*_*s*_, sin *θ*_*s*_) during either the response period (delay task) or the stimulus period (react task); and is changed to (cos(*θ*_*s*_ +*π*), sin(*θ*_*s*_ +*π*)) for their anti versions. For the match-to-sample tasks, the target is (*y*_1_, *y*_2_) = (cos *θ*_*m*_, sin *θ*_*m*_) during the response period if *θ*_*m*_ = *θ*_*s*_, and 0 otherwise as well as during other periods; for the anti version, the condition is changed to *θ*_*m*_ = *θ*_*s*_ + *π* (Fig. 7A1).

#### Categorization tasks

This family comprises four tasks, parameterized by two constant task inputs: *c*_in_ ∈ {±1}, selecting the relevant stimulus channel (*c*_in_ = +1 for *s*_1_, *c*_in_ = −1 for *s*_2_), and *c*_out_ ∈ {±1}, selecting the output channel reporting the categorization result (*c*_out_ = +1 for *y*_1_, *c*_out_ = −1 for *y*_2_), inspired by the experimental design of Tafazoli et al. [16] (Fig. 6A). The input is five-dimensional, *s* = (*s*_1_, *s*_2_, *f, c*_in_, *c*_out_), where *s*_1_, *s*_2_ ∈ [−1, 1] encode two distinct stimulus features, and *f* ∈ {0, 1} is the fixation input. The target is two-dimensional, *y* = (*y*_1_, *y*_2_) ∈ ℝ^2^. As in the previous ‘memory’ tasks, there are four consecutive periods: context, stimulus, memory, and response.

Task input is constant across all periods. During the context period, the stimulus is zero and *f* = 1. During the stimulus period, *s*_1_ and *s*_2_ take non-zero values and *f* = 1. During the memory period, all stimulus inputs are zero and *f* = 1. During the response period, all stimulus inputs are zero and *f* = 0. The network must categorize the relevant stimulus, 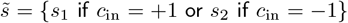, as positive or negative, and report the result in the selected output channel. The target output during the response period is therefore 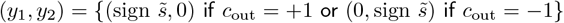, and 0 during other periods.

### E. Task solutions

#### Memory pro / anti tasks (shared)

The latent space is two-dimensional, *λ* = (*λ*_1_, *λ*_2_) ∈ ℝ^2^ and the readout is two-dimensional, *y* = (*y*_1_, *y*_2_) ∈ ℝ^2^. The full input is *s* = (*s*_1_, *s*_2_, *f, c*) where (*s*_1_, *s*_2_) ∈ ℝ^2^ are the stimulus input, *f* ∈ {0, 1} the fixation input, and *c* ∈ {−1, 1] encodes the pro (*c* = 1) or anti (*c* = −1) version of the tasks. We use polar coordinates for (*λ*_1_, *λ*_2_) and (*s*_1_, *s*_2_) with radii *r, r*_*s*_ and angles *θ, θ*_*s*_, respectively.

#### Flow field

The autonomous radial dynamics in the latent space are given by a quartic double-well potential with stable fixed points at *r* = 0 and *r* = 1, corresponding to the radial flow

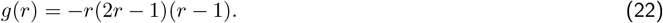

The total autonomous flow field is given by

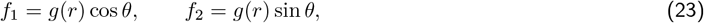

implementing a fixed point at the origin and a ring attractor. The input-driven flow field is

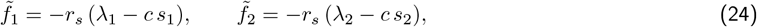

which leads to a fixed point at *λ*_1_ = *c s*_1_, *λ*_2_ = *c s*_2_ where the multiplication with *c* rotates the angle by *π* for the anti tasks. The multiplication with *r*_*s*_ ensures that there are no input-driven dynamics without a stimulus input. The readout is

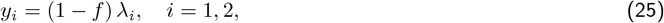

reporting the latent state if fixation is switched off.

#### Training distribution

The stimulus (*s*_1_, *s*_2_) = *r*_*s*_(cos *θ*_*s*_, sin *θ*_*s*_) is sampled according to *θ*_*s*_ ~ Unif[0, 2*π*] and *r*_*s*_ ~ Bernoulli(*c* = 0.25). Further, *f* ~ Bernoulli(*c* = 0.5) and *c* ~ Rademacher({−1, +1}). The latent state is sampled as *λ* ~ N(0, *σ*^2^*I*_2_) with *σ* = 0.5. With probability *p*_norm_ = 0.25 the planar part is normalized, 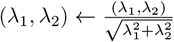, if there is no stimulus input to increase the number of samples on the ring attractor.

#### Memory pro / anti tasks (non-shared)

We use the same notation as in the shared solution above, and specify only differences.

#### Parallel solution

For the “parallel” example, the latent space is three-dimensional and the autonomous flow-field is task-dependent: the flow field is identical to the shared solution in the (*λ*_1_, *λ*_2_)-plane, but the flow-field along the *λ*_3_-axis depends on the task input *c*: *f*_3_ = − (*λ*_3_ −*c*), such that the ring attractor is in the *λ*_3_ = *c* plane. The input-driven flow field is task-invariant: the multiplication with the task input *c* in Eq. (24) is replaced by a multiplication with *λ*_3_, 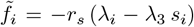 for *i* = 1, 2; there is input-driven flow in the *λ*_3_ direction. The readout is identical to the shared solution, and does not depend on *λ*_3_. The training data distribution is identical except that *λ* is drawn from a three-dimensional normal distribution, *λ*~N (0, *σ*^2^*I*_3_); and when normalization occurs, it is on the ring in the plane *λ*_3_ = *c*.

#### Reflected solution

For the “reflected” example, the latent space is two-dimensional and the autonomous flow-field is task-dependent. The full latent space is shifted by (1, 0) for the pro task, and then reflected with respect to the *y*-axis for the anti task. Within the shifted latent space, the input-driven dynamics are identical to that of the shared solution — for the anti task, input-driven dynamics are the reflected opposite of that of the pro task. The readout is task-invariant: it reports the latent state, projected on the (*λ*_1_ ≥ 0)-half-plane, and shifted by (1, 0), i.e. ***y*** = (1 − *f*) · (|*λ*_1_| − 1, *λ*_2_). The training data distribution is also identical, except that the planar normalization is also shifted and reflected according to the task to match the location of the ring attractor.

#### Orthogonal solution

For the “orthogonal” example, the latent space is four-dimensional and the autonomous flow-field is task dependent: for the ‘pro’ task, the flow field is identical to the shared solution in the (*λ*_1_, *λ*_2_)-subspace and the autonomous flow in the (*λ*_3_, *λ*_4_)-subspace is contractive, *f*_3_ = −*λ*_3_ and *f*_4_ = −*λ*_4_; for the ‘anti’ task, the flow field is identical to the shared solution in the (*λ*_3_, *λ*_4_)-subspace and the autonomous flow in the (*λ*_1_, *λ*_2_)-subspace is contractive. The input flow field is task-invariant: 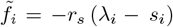 for *i* = 1, 2 and 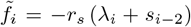 for *i* = 3, 4, such that input-driven dynamics solve the pro task in the (*λ*_1_, *λ*_2_)-subspace and the anti task in the (*λ*_3_, *λ*_4_)-subspace. The readout flow field is also task-invariant: *y*_*i*_ = (1 − *f*) (*λ*_*i*_ + *λ*_*i*+2_) for *i* = 1, 2, leveraging that the flow is contractive in the irrelevant subspace. The training data distribution is identical except that *λ* is drawn from a four-dimensional normal distribution, *λ* ~ N(0, *σ*^2^*I*_4_) and planar normalization is performed independently in the two subspaces (*λ*_1_, *λ*_2_) and (*λ*_3_, *λ*_4_).

#### Ring attractor family

We use the same notation as in the shared memory task above, and specify only differences. The latent space is three-dimensional, *λ* = (*λ*_1_, *λ*_2_, *λ*_3_) ∈ ℝ^3^ and the full input is *s* = (*s*_1_, *s*_2_, *f, c*_0_, *c*_1_, *c*_2_) where *c*_1_ 0, 1 encodes whether the match-to-sample (*c*_1_ = 1) or another task (*c*_1_ = 0) should be solved, *c*_2_ ∈ {0, 1} distinguishes react (*c*_2_ = 1) from delay and memory tasks (*c*_2_ = 0), and *c*_0_ encodes the pro/anti context (as *c* in the memory tasks above).

#### Flow field

The autonomous radial dynamics in the (*λ*_1_, *λ*_2_)-plane are given by the fixed point and the ring attractor, and the flow along the *λ*_3_ axis is contractive towards *λ*_3_ = *r* with 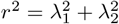. In total, the autonomous flow field is

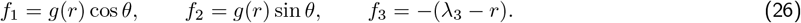

The flow in the *λ*_3_ tracks the radius in the (*λ*_1_, *λ*_2_)-plane; since the ring has radius *r* = 1 this shifts the ring into the *λ*_3_ = 1 plane.

The input-driven flow field is

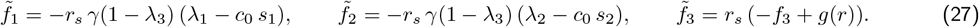

where 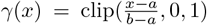 with *a* = 0.1, *b* = 0.9 interpolates between *γ*(*x* ≤ *a*) = 0 and *γ*(*x* ≥ *b*) = 1. The multiplication with *γ*(1 − *λ*_3_) gates the input such that it has an effect in the *λ*_3_= 0 plane but not in the *λ*_3_= 1 plane. The double-well along the *λ*_3_ direction ensures that the dynamics take place in the *λ*_3_ = 0 plane during the first stimulus presentation.

The output is

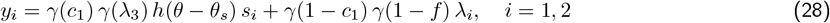

where *h*(*θ*) = exp[*ν*(cos *θ* − 1)] with *ν* = 5. Here, *γ*(*c*_1_) and *γ*(1 − *c*_1_) gate the output to the term for the match-to-sample or the memory task, respectively. The first term, corresponding to the output for match-to-sample tasks, is only non-zero if the network is on the ring attractor in the *λ*_3_ = 1 plane due to the factor *γ*(*λ*_3_), that is a first stimulus has been presented, and the angle of the current stimulus is close to the angle on the ring attractor due to the factor *h*(*θ* −*θ*_*s*_). If both conditions are fulfilled, the output is the strength of the stimulus. The second term, corresponding to the memory tasks, is non-zero if the fixation is off, implemented by the factor *γ*(1 − *f*), and reports the latent state in the (*λ*_1_, *λ*_2_)-plane, i.e., on the ring once if a stimulus was presented. Finally, to solve the ‘react’ tasks, the input weights for *c*_2_ were set equal to the negative input weights from the fixation input *f*.

#### Training distribution

The stimulus (*s*_1_, *s*_2_) = *r*_*s*_(cos *θ*_*s*_, sin *θ*_*s*_) is sampled according to *θ*_*s*_ ~ Unif[0, 2*π*] and *r*_*s*_ ~ Bernoulli(*p*_stim_) with *p*_stim_ = 0.25. Further, *f* ~ Bernoulli(*p*_fix_), 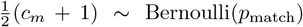, and *c*_0_ ~ Rademacher({−1, +1}), with *p*_match_ = 0.4 and *p*_fix_ = 0.5. The latent state is sampled as *λ* ~ N(0, *σ*^2^*I*_3_) with *σ* = 0.5. With probability *p*_norm_ = 0.2 the planar part is normalized, 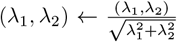, and independently *λ*_3_ is fixed to *λ*_3_ = 1 with probability *p*_clip*z*_ = 0.2, to increase the number of samples on the ring attractor.

#### Categorization tasks (re-used across all tasks)

We use the same notation as in the shared memory task above, and specify only differences. The latent space is one-dimensional, *λ*_1_ ∈ ℝ and the full input is *s* = (*s*_1_, *s*_2_, *f, c*_in_, *c*_out_) where *c*_in_ ∈ {−1, 1} selects the input channel, *c*_in_ = 1 for *s*_1_ and *c*_in_ = −1 for *s*_2_, and *c*_out_ ∈ {−1, 1} selects the output channel, *c*_out_ = 1 for *y*_1_ and *c*_out_ = −1 for *y*_2_.

#### Flow field

The autonomous flow field implements stable fixed points at *λ*_1_ = ±1 and *λ*_1_ = 0, 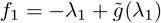 with

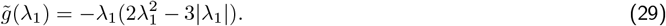

The input-driven flow field is

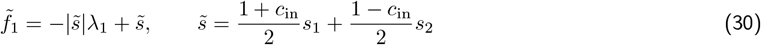

driving the dynamics towards the fixed point 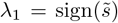 according to the relevant input channel selected via 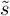. The output is

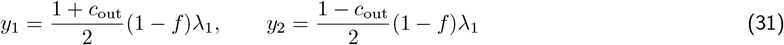

which reports the latent state in the relevant output channel.

#### Training distribution

The stimuli 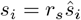 are sampled according to 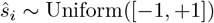 and *r*_*s*_ ∼ Bernoulli(*p* = 0.25). Further, *f* = 1 if *r*_*s*_ = 1 and otherwise *f* ∼ Bernoulli(*p* = 0.5), *c*_in_ ∼ Rademacher(1, +1), and *c*_out_ ∼ Rademacher(1, +1). The latent state is sampled as *λ*_1_ (0, *σ*^2^) with *σ* = 0.5. With probability *p*_norm_ = 0.25 the latent space is normalized, 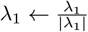, if there is no stimulus input to increase the number of samples on the fixed points.

#### Categorization tasks (re-used across some tasks)

We use the same notation as in the re-used solution for the categorization task, and specify only differences. The latent space is two-dimensional, ***λ*** = (*λ*_1_, *λ*_2_) ∈ ℝ^2^, while inputs and outputs remain identical.

#### Fully invariant solution

For the fully invariant solution, the autonomous flow field is

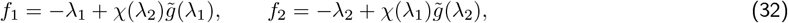

where *χ*(*λ*) = 1 − min(1, |*λ*|) is a ‘tent-shaped’ gate with *χ*(0) = 1 and *χ*(±1) = 0, such that there are five attractors at (0, 0), (0, 1), (0, −1), (1, 0), and (−1, 0). The input-driven flow field is

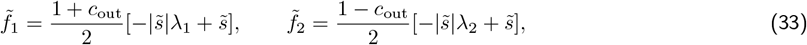

which is identical to the re-used solution except that it resolves the output channel. This allows for a final output that is shared across all tasks, *y*_*i*_ = (1 − *f*)*λ*_*i*_ for *i* = 1, 2.

#### Partially invariant solution

For the partially invariant solution, the autonomous flow field depends on the task input:

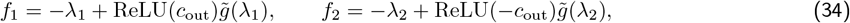

such that there are only attractors along the *λ*_1_-axis if *c*_out_ = −1 and along the *λ*_2_-axis if *c*_out_ = − 1. The input-driven flow field is

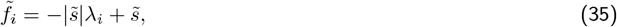

driving the dynamics to 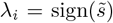 independent of *c*_out_, but the autonomous dynamics lead back to the origin for the irrelevant output dimension. This allows again for a final output that is shared across all tasks, *y*_*i*_ = (1 − *f*)*λ*_*i*_ for *i* = 1, 2.

#### Training distribution

The full input is distributed as in the re-used solution. The latent state is sampled as ***λ*** ~ N(0, *σ*^2^*I*_2_) with *σ* = 0.5. With probability *p*_fp_ = 0.25 the latent state is set to an attractor chosen with equal probability from {(0, 1), (0, −1), (1, 0), (−1, 0)} to increase the number of samples on the fixed points.

### F. A family of solutions to the two-factors categorization tasks

Each of the solutions to the categorization tasks presented in Fig. 6 is characterized by the structure of attractor re-use and the loci of task-dependence — i.e., which attractors are re-used across which task conditions, and where each of the two contextual factors (***c***_in_, ***c***_out_) enters the computation, as summarized in Fig. 6E. Building on this systematic characterization, we extended the set of solutions to build a family of nine qualitatively different solutions (Fig. S1). Importantly, we only consider solutions where the irrelevant stimulus is not persistently encoded via attractor dynamics, consistent with PFC data [16, 17].

**Figure S1.**
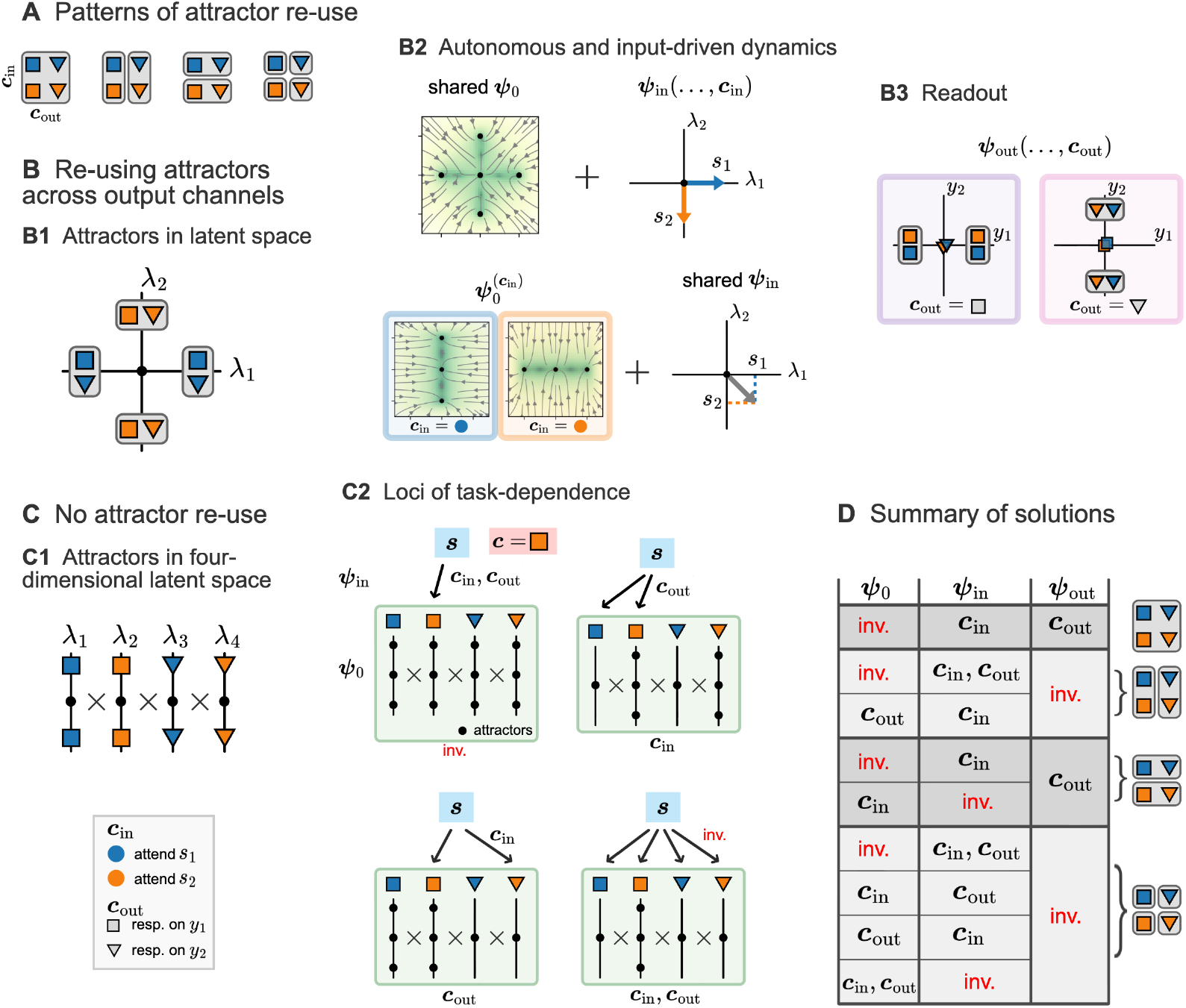
Nine qualitatively different solutions to the categorization tasks of Fig. 6. **A**. Four possible patterns of attractor re-use across the four task conditions, (***c***_in_, ***c***_out_) ∈ {±1} ^2^. **B**. Solutions with attractors re-used across tasks with the same ***c***_out_. **B1**. Latent attractors in a two-dimensional space; one pair of attractors per value of ***c***_out_. **B2**. Two solutions distinguished by the locus of ***c***_in_-dependence: ***ψ***_in_ (top) or ***ψ***_0_ (bottom); the complementary component is ***c***_in_-invariant. Plotting conventions as in Fig. 6. **B3**. Readout ***ψ***_out_, modulated by ***c***_out_. Markers represent ***ψ***_out_ evaluated at the attractors of panel (B1), for the two values of ***c***_out_. **C**. Solutions with no attractor re-use. **C1**. Latent attractors in a four-dimensional space, with one independent pair in an orthogonal latent dimension for each task condition. **C2**. Four solutions, each characterized by a distinct combination of loci through which ***c***_in_ and ***c***_out_ enter the computation: ***ψ***_in_ (arrows) or ***ψ***_0_ (boxes), with ***c***_in_ and ***c***_out_-dependence indicated. For an example task ***c*** (orange square: attend *s*_2_, respond on *y*_1_), arrows represent the task-dependent routing of the input ***s*** through ***ψ***_in_ towards specific latent dimensions (*λ*_1,2,3,4_); and dots on lines in green boxes represent the task-dependent existence of attractor pairs along each latent dimension. Since attractors are different across response directions ***c***_out_, the readout ***ψ***_out_ need not be task-dependent. **D**. Summary of all nine solutions. Rows represent solutions, grouped by attractor re-use pattern (right); columns show the dependence of ***ψ***_0_, ***ψ***_in_, and ***ψ***_out_ on ***c***_in_ and ***c***_out_ (‘inv.’: task-invariant). First three rows are shown in Fig. 6.

For each of the four categorization tasks, the network needs a pair of attractors to store the categorization result. In addition to the cases where attractors were re-used across all tasks (Fig. 6B) and only across tasks with the same response direction (***c***_out_; Fig. 6C), attractors can be shared only across tasks with the same relevant input feature (***c***_in_); finally, the network could use a different pair of attractors for each task. This leads to four distinct patterns of attractor re-use (Fig. S1A).

When several task subsets use distinct attractors, there are two possibilities: all attractors can be constantly present in the autonomous dynamics, so that task-dependence must be resolved by the input dynamics ***ψ***_in_ (as in Fig. 6C2); alternatively, task-dependence can modulate the autonomous dynamics (as in Fig. 6C3), so that only attractors relevant to the correct task subset are present, and input dynamics need not be task-dependent. This reasoning also holds when attractors are shared only across ***c***_out_, yielding two distinct solutions (Fig. S1B). Finally, for both solutions, ***c***_out_-dependence must be resolved by the readout ***ψ***_out_ (Fig. S1B3).

In the case where a different pair of attractors is used for each task, both contextual factors (***c***_in_, ***c***_out_) need to be resolved either by the autonomous dynamics 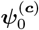 or the input ***ψ***_in_; while the readout ***ψ***_out_ can be task-invariant. The four possible combinations of factors and loci lead to four distinct solutions (Fig. S1C), each characterized by specific invariances of the autonomous and input-driven dynamics. This leads to a systematic catalogue of nine qualitatively different solutions, each uniquely characterized by its pattern of attractor re-use and the loci at which ***c***_in_ and ***c***_out_ modulate the computation (Fig. S1D). Note that we did not consider solutions with additional unnecessary task dependence; for instance, a solution where both ***ψ***_in_ and ***ψ***_0_ would resolve ***c***_out_.

## Footnotes

1 The low-pass filter is 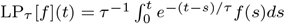.

## Notes

### Competing Interest Statement

The authors have declared no competing interest.

